# The *Legionella pneumophila* type IVb secretion system effector BinA subverts amino acid transport to sensitize TORC1 signaling in macrophages

**DOI:** 10.1101/2025.02.20.639245

**Authors:** Magdalena Circu, Reneau Castore, Brian Latimer, Stephanie Shames, Craig R. Roy, Ana-Maria Dragoi, Stanimir S. Ivanov

**Affiliations:** Department of Microbiology and Immunology, Louisiana State University Health Sciences Center - Shreveport, Shreveport, Louisiana, USA; Department of Molecular and Cellular Physiology, Louisiana State University Health Sciences Center - Shreveport, Shreveport, Louisiana, USA; Innovative North Louisiana Experimental Therapeutics program (INLET), Feist-Weiller Cancer Center, Louisiana State University Health Sciences Center - Shreveport, Shreveport, Louisiana, USA; Department of Microbiology, Genetics, and Immunology, Michigan State University, East Lansing, Michigan, USA; Department of Microbial Pathogenesis, Yale University School of Medicine, 295 Congress Avenue, New Haven, Connecticut, USA

**Author notes:** corresponding author, (SSI). Department of Pathology, Louisiana State University Health Sciences Center - Shreveport, Shreveport, Louisiana, USA.

## Abstract

*Legionella pneumophila* is an environmental Gram-negative bacterium that parasitizes unicellular protozoa and can cause severe pulmonary infections when aerosolized bacteria are inhaled by humans. One critical aspect of *Legionella* pathogenesis is the establishment in the cytosol of infected macrophages of a unique ER-derived vacuole, that requires a sustained supply of host lipids during expansion. Subversion of pro-lipogenic pathways downstream of the metabolic checkpoint kinase mTOR (Mechanistic Target of Rapamycin) are critical for niche expansion. In eukaryotic cells, amino acids sufficiency and growth factor sensory signals converge on mTOR to ensure metabolic processes are coupled to nutrients/energy availability. *Legionella* can trigger mTOR signaling in infected cells by increasing the intracellular abundance of amino acids through inhibition of host translation. Here, we describe a novel mechanism by which *Legionella* sensitizes mTOR in infected macrophages. A forward genetic screen identified Lpg0393 protein as a putative bacterial mTOR regulator that contains a VPS9-domain typically found in eukaryotic GEFs (Guanine nucleotide exchange factors) for Rab5 GTPase family members (Rab5/Rab21/Rab22). We uncovered that Lpg0393 lowers the activation threshold for mTOR signaling upon stimulation with arginine or leucine through rewiring of processes upstream of mTOR by removing Rab5-dependency and replacing it with a Rab21/Rab22-dependency. Data from cells expressing either a bacterial or a eukaryotic mTOR sensitizing factor uncovered two distinct intraorganellar Arg/Leu pools that fuel mTOR activation in parallel – one regulated by Rab21/22 and the other by Rab5. Consistent with the role of mTOR in expansion of the *Legionella*-occupied organelle, deletion of Lpg0393 also resulted in premature vacuolar rupture in a mTOR-dependent manner. All together, we have identified a novel bacterial mTOR regulator and consistent with its reported functions we propose Lpg0393 is named as BinA (**<u>B</u>**acterial **<u>in</u>**itiator of TORC1 signaling and an **<u>a</u>**ctivator of Rab5 family GTPases).

**Author Summary:** *Legionella pneumophila* - a prototypical vacuolar pathogen – manipulates host lipogenesis to sustain membrane biogenesis, which ensures the integrity of the vacuolar compartment is not compromised as demand for housing capacity increases during bacterial replication. Subversion of the host mTOR kinase signaling, which functions as a central regulatory hub coordinating nutrients sufficiency with metabolic output, is one mechanism by which *Legionella* increases *de novo* lipogenesis in infected cells. Here, we report a novel mechanism by which the bacteria sustain mTOR signaling through the actions of the type IV secretion system effector BinA. We found that BinA rewires amino acid transport mechanisms by manipulating Rab5 family GTPase to selectively sensitize mTOR to arginine and leucine but not methionine or glutamine stimulation. mTOR sensitization by BinA revealed a switch in upstream regulators from Rab5 to Rab21/Rab22. Our work provides novel insight into how pathogens exert metabolic control over host cells to maximize intracellular replication.

## Introduction

*De novo* biogenesis and maintenance of a unique membrane-bound organelle is a common challenge faced by intracellular pathogens that is frequently resolved through subversion of certain aspects of host metabolism [1–4]. One example is the exploitation of host lipogenesis by the intravacuolar bacterial pathogen *Legionella pneumophila* through the activation of the host metabolic coordinator mTOR [4–6]. The metabolic checkpoint serine/threonine kinase mTOR orchestrates synchronization of anabolic and catabolic processes in eukaryotic cells and thus is wired to multiple nutrients sufficiency response pathways, including growth factors and amino acid abundance, to ensure biosynthetic processes are coupled with precursors availability [7–12]. *De novo* lipogenesis is one mTOR-regulated anabolic process producing phospholipids and cholesterol from metabolic precursors at the ER for the biogenesis of cellular membranes [13–15].

mTOR functions as part of two kinase complexes known as TORC1 and TORC2, where each complex has a distinct composition (although some components are shared), distinct substrate specificities and distinct subcellular localizations [12, 16–18]. Raptor and Rictor are defining scaffolding proteins for TORC1 and TORC2 respectively that dictate substrate specificity [12, 18]. Pharmacological and genetic studies attribute a central role for TORC1 in *de novo* lipogenesis in eukaryotic cells [10, 13, 19]. Signals from amino acid abundance and growth factor availability sensors are transmitted through a complex layered regulatory network and merge at the lysosomal membrane to activate TORC1 [20, 21]. Various amino acid-binding proteins monitor sufficiency and act upstream of the heterodimeric Rag GTPase complex, which functions as a molecular switch and depending on its nucleotide-bound state recruits TORC1 to lysosomal membranes for activation under amino acids replete conditions [20]. This process is mediated in part by the heteropentameric Ragulator complex, composed of the LAMTOR1/2/3/4/5 subunits, which tethers the Rag GTPase complex to the lysosomal membrane and promotes GTP release from RagC, thus removing one barrier to Rag GTPase complex activation [22–24]. Once localized to the lysosomal membrane, TORC1 is directly activated by the small GTPase Rheb, which acts downstream of the growth factor signaling axis PI3K-Akt-TSC2 [25].

*Legionella pneumophila (L.p)* is an accidental human pathogen that causes a severe lower respiratory infection known as Legionnaires’ disease [26, 27], where alveolar macrophages represent a major cellular reservoir that supports bacterial replication [28–30]. As a facultative intracellular bacteria, *L.p* replicates intracellularly within a unique ER-derived vacuolar organelle referred to as the LCV (*Legionella*-containing vacuole) in its natural host unicellular protozoa as well as in mammalian macrophages [31–35]. Macrophages can also support extracellular *L.p* replication under conditions of compromised nutritional immunity [36]. The molecular survival strategies of *L.p* which evolved through adaptation to a broad range of protozoan facilitate colonization and replication within mammalian macrophages [37]. Upon uptake, *L.p* blocks endocytic maturation and initiates phagosomal membrane remodeling through the recruitment and fusion with early secretory vesicles [32, 38, 39] and ER-derived membranes [40] to initiate LCV biogenesis. Mature remodeled LCVs support bacterial replication starting at ∼ 4 hours post uptake, however differences in LCV lifespan can produce high variability in the number of bacteria released after LCV rupture [4, 41–43]. LCV lifespan and housing capacity has been linked to mTOR activation as well as initiation of *de novo* lipogenesis in *L.p* infected macrophages; thus, inhibition of these processes results in premature LCV rupture [4, 5, 44]. *L.p* infection triggers sustained TORC1 signaling, even in the absence of growth factor stimulation, specifically in macrophages harboring bacteria but not in bystander uninfected cells [4, 6]. This phenomenon requires a functional type IVb secretion system (T4bSS), known as the Dot/Icm apparatus that translocates over 300 bacterial effector proteins directly into the host cytosol [45–47]. *L.p*-induced increase of cytosolic amino acids in infected macrophages is proposed to sustain TORC1 signaling [48]. The host translation machinery is targeted by a number of T4bSS effectors that block protein synthesis, including the glycosyltransferases Lgt1, Lgt2 and Lgt3 [49, 50], which in turn stops amino acid consumption by the translation machinery effectively increasing the abundance of cytosolic amino acid leading to TORC1 activation [48]. Similar to *L.p* infections, pharmacological inhibitors targeting the protein initiation or elongation machinery also trigger TORC1 signaling in the absence of growth factor stimulation [51]. However, the current model only partially explains the experimental data. TORC1 activity is triggered within 3hpi and last for the duration of the LCV lifespan when infections are carried out in amino acid-containing medium [4]. *L.p*-driven TORC1 signaling can tolerate withdrawal of exogenous amino acids for short duration (∼60 min), in a manner requiring host-translation blocking *L.p* effectors [48]. However, bacteria-induced TORC1 signaling is completely inhibited upon prolonged 4-hour withdrawal of exogenous amino acids [4] indicating that additional mechanisms allow *L.p* to sustain TORC1 signaling in host cells. In this study, we discovered that *L.p*-induced TORC1 signaling in infected macrophages is sustained by continuous import of amino acids from the extracellular/intraorganellar pools and this process is subverted via the *L.p* T4bSS effector BinA through manipulation of host Rab GTPases that regulate early endocytic membrane trafficking.

## Results

### Forward genetic screen for *L.p*-encoded TORC1-activating factors identified the T4bSS effector BinA as a putative regulator

One mechanism mediating bacteria-induced TORC1 activation in *L.p* infected macrophages is driven by the accumulation of intracellular amino acids caused by a *L.p* - induced blockade in host protein translation [48] (Fig 1A). Previous data also supports the idea of T4bSS effectors-mediated TORC1 activation [4, 48], thus we carried out an immunofluorescence-based forward genetic screen to identify putative bacterial mTOR regulators using a cellular model of infection (Fig 1A-C). To this end, *Myd88*^-/-^ bone marrow-derived macrophages (BMDMs) were selected as host cell model because previous data demonstrates that under these conditions TORC1 activation (i) is confined to cells harboring bacteria and (ii) is triggered solely by *L.p* in a T4SS-dependent manner [4]. Thus, putative bacterial inducers of mTOR signaling can be identified in this setting.

**Fig 1.**
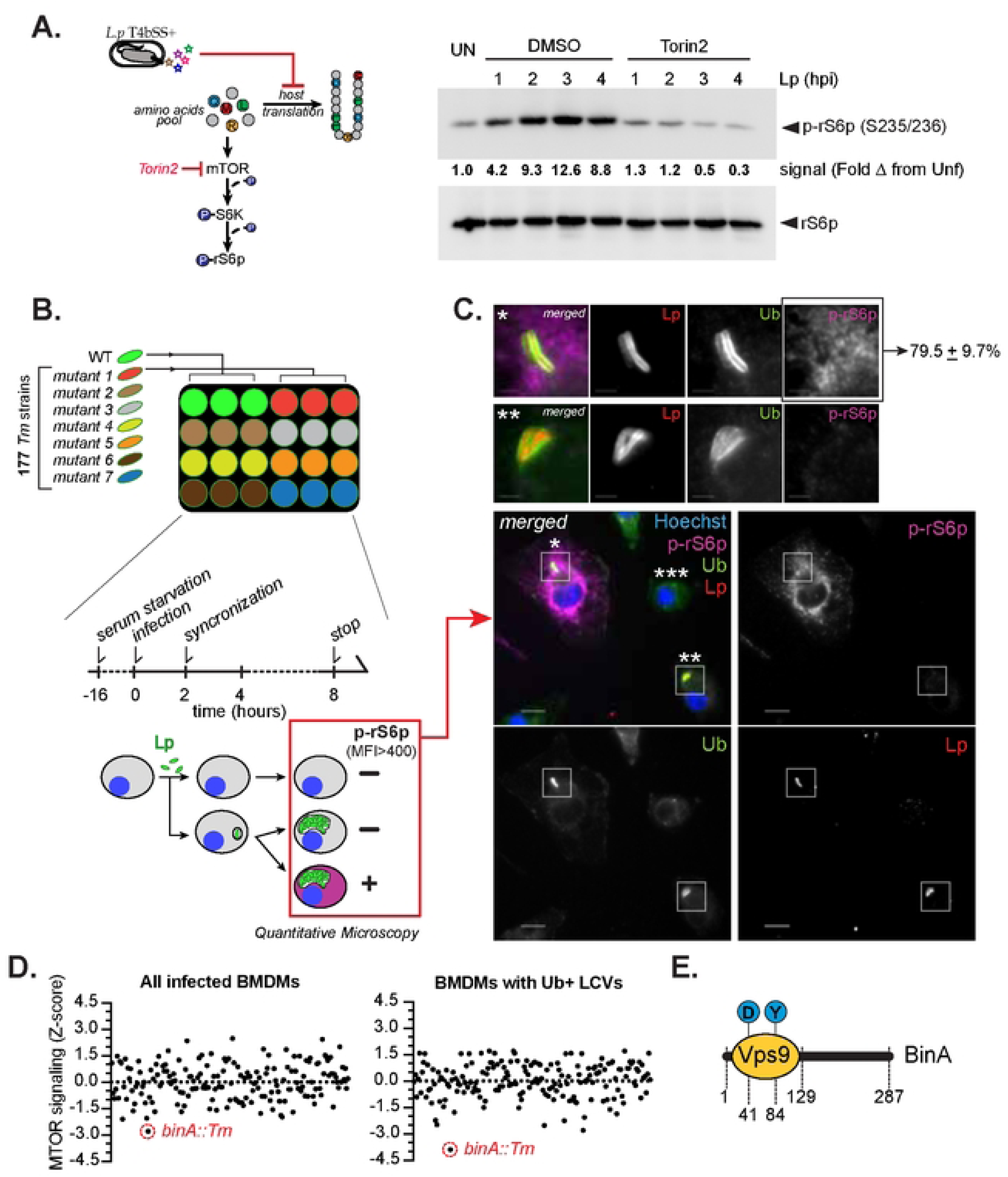
Identification of Lp T4bSS effector regulating TORC1 signaling. **(A)** TORC1 signaling induced by Lp results in phosphorylation of the host ribosomal S6 protein (rS6p). (A, left panel) Lp can activate TORC1 by suppressing host translation, which increases the intracellular amino acids concentration [4, 48]. (A, right panel) Immunoblot (IB) analysis showing rS6p phosphorylation at Ser235/236 in cell lysates from Myd88 KO iBMDMs infected with Lp01 Δ*flaA* (MOI=15) for the indicated time periods in the presence or absence of the MTOR inhibitor Torin2 or were left uninfected (UN). **(B)** Schematic depiction of the performed loss-of-function screen for TORC1 activators with an arrayed library of single-gene transposon mutants strains lacking individual T4bSS effectors. Myd88 KO BMDMs were seeded on glass cover slips and were infected for 8hrs, one strain at a time where each plate included Lp01 Δ*flaA* baseline infection (MOI=15). Cover slips were immunostained for p-rS6p (S235/236) and ubiquitin followed by quantitative microscopy analysis of the percentage of infected p-rS6p+ macrophages. **(C)** Representative micrograph of Lp01 Δ*flaA* infection showing (*) an infected p-rS6p+ cell, (**) an infected p-rS6p− cell, and (***) an uninfected p-rS6p− cell. The average baseline percentage of p-rS6p+ cells in Lp01 Δ*flaA* infections was 79.5 ± 9.7%. **(D)** Data shows the distribution of the standard deviations away from the mean (z-score) for each mutant strain tested. Data is parsed out for macrophages housing remodeled Ub+ LCVs as well as for all infected cells. The results for the top hit *binA*::Tm are highlighted in both categories. The complete dataset is provided in Table S1. **(E)** Domain architecture of BinA is shown. Residues D41 and Y84 are critical for the GEF activity of the VPS9-like domain.

Phosphorylation of the ribosomal S6 protein (rS6p) at Ser235/236 by the S6 protein kinase (S6K) is a canonical downstream readout for TORC1 activation, which was incorporated in the screen. In *Legionella* infection assays, phospho-rS6p abundance can be monitored via immunoblot analysis at the population level (Fig 1A) [4] and via immunofluorescence microscopy at the single cell level (Fig 1C). We screened an arrayed T4SS effectors library of 177 single gene transposon mutants [52] in synchronized cellular infection assays lasting 8 hours, where each infection was carried out in a technical triplicate (Fig 1B). The p-rS6p signal in each infected cell was determined via immunofluorescence where ‘ON’ state was set at p-rS6p mean fluorescent signal intensity > 400 based on data from bystander uninfected cells as well as cells infected by a mutant strain lacking a functional T4SS – *L.p* Δ*dotA* (Fig 1C) [4]. Because deletion of T4SS effectors might produce intracellular colonization defects due to failure in LCV trafficking or remodeling, immunostaining with anti-Ubiquitin (FK2) antibody was also included in the analysis so that LCV defects can be measured as an experimental variable. Ubiquitin accumulation at the LCV coincides with initiation of LCV biogenesis and persists for the lifespan of the vacuole [53]. Representative BMDMs with distinct TORC1 signaling states (as defined for the screen) that harbor remodeled Ub+ LCVs as well as uninfected bystander cells are shown in Fig 1C. For each set of experiments, infections with wild-type *L.p* were included as a baseline for the percentage of infected cells that trigger rS6p phosphorylation, which was determined to be 79.5 ± 9.7% (Fig 1C). Two z-scores were calculated for each mutant strain taking into account all infected cells or only BMDMs harboring Ub+ LCVs, which allowed for identification of false positives due to intracellular colonization defects (Fig 1D). The top hit in both categories was a mutant strain with an insertion in the *binA* gene (*lpg0393*) (Fig 1D-E). BinA has been shown to be a *bona fide* substrate of the *Legionella* Dot/Icm T4bSS [54] that contains a eukaryotic VPS9 domain typically found in GEFs that activate Rab5 family of small GTPases [55]. BinA exhibits GEF activity *in vitro* towards Rab5, Rab21 and Rab22, where D41 and Y84 represent key catalytic residues in the VPS9 domain required for nucleotide exchange [56]. The role of BinA in *Legionella* infections remains to be elucidated.

To investigate BinA as a putative TORC1 regulator we generated a *binA* clean deletion strain, which was subsequently complemented with a plasmid-borne 3xFLAG- tagged copy of *binA* under the control of an IPTG-inducible promoter (Fig 2A). The capacity of those strains to trigger rS6p phosphorylation in *Myd88*^-/-^ BMDMs was assessed at distinct time-points after infection (Fig 2B-D). To this end, the p-rS6p signal intensity was measured in synchronized infections at the single-cell level via immunofluorescence where time points were selected to capture TORC1 signaling at the time of initiation of bacterial replication (4hpi), after 1-2 doubling events (6hpi) or after several rounds of replication (10hpi) have taken place (Fig 2B-D). Macrophages harboring *binA–* bacteria across all time point produced significantly lower p-rS6p signal at the population level as compared to BMDMs infected by the wild-type or the complemented strains (Fig 2D). The p-rS6p signal in some cells infected by *binA–* bacteria was comparable to cells harboring *binA*+ strains, however in majority of the cell population the signal was lower indicating that either BinA is required for optimal TORC1 signaling or functional redundancy from other bacterial factors partially compensates for the loss of *binA*. The percentage of Ub+ LCVs harboring *binA–* bacteria at 6hpi was comparable to LCVs containing *binA*+ bacteria (82.4 ± 7% vs 77.8 ± 9%) indicating that loss of *binA* did not produce LCV maturation or trafficking defects. Importantly, the induction of p-rS6p was restored in macrophages infected by the *binA* complemented strain under IPTG-inducing conditions to a level indistinguishable from the wild type parental strain (Fig 2B-D). Taken together, these data indicate a role for BinA as a putative bacterial TORC1 regulatory factor.

**Fig 2.**
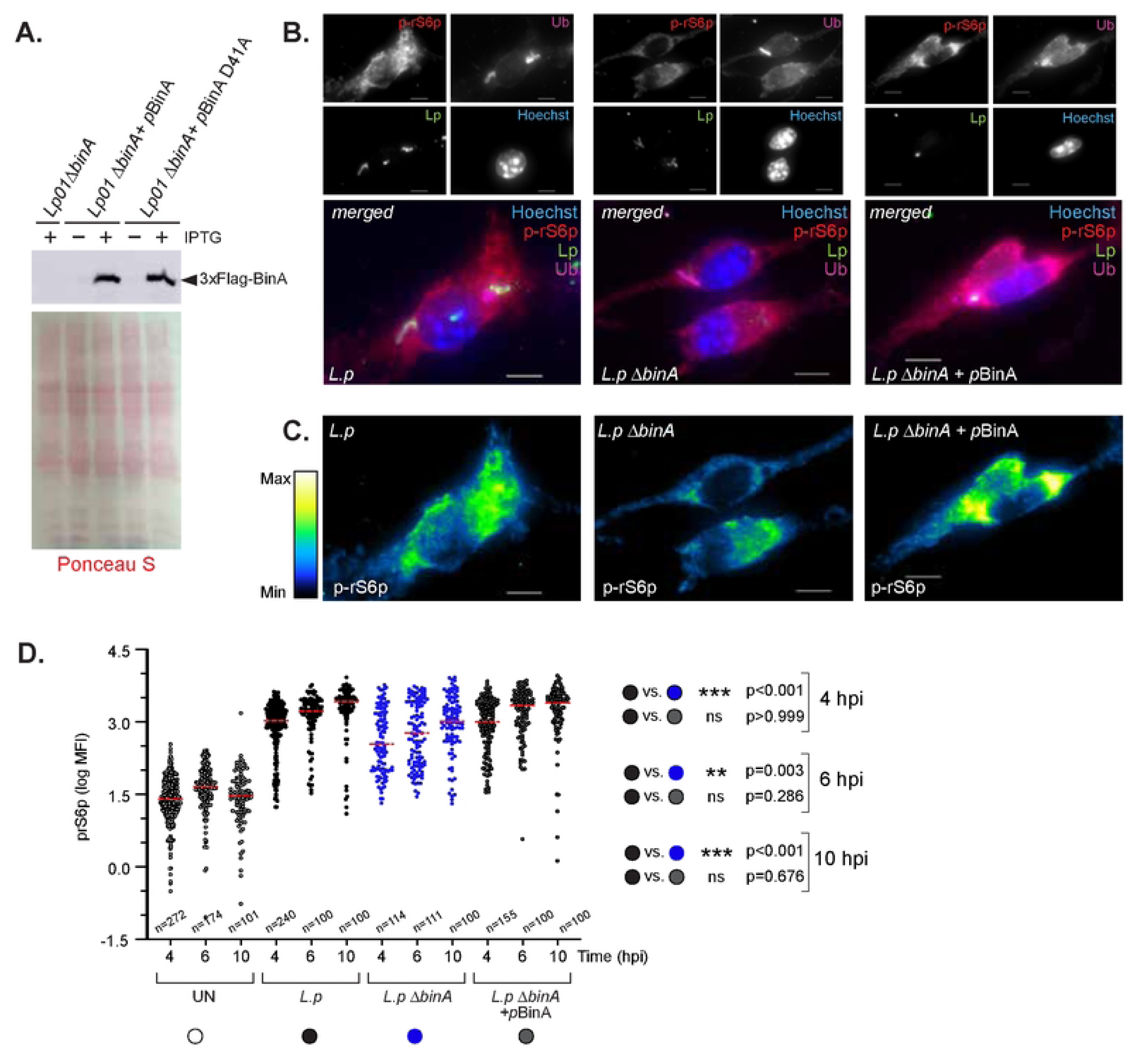
Deletion of BinA compromises TORC1 activation by Lp. **(A)** Anti-Flag immunoblot analysis of various Lp strains expressing the indicated plasmid borne IPTG- inducible 3XFlag-BinA alleles. Ponceau S staining is used to control for lysate loading. **(B)** Representative micrographs of TORC1 activation in Myd88KO iBMDMs infected by Lp strains lacking or expressing BinA at 6hpi. Phosphorylation of rS6p was used as a readout for TORC1 signaling and is shown as pseudocolor in **(C)**. **(D)** Quantitative microscopy of the p-rS6p signal from uninfected (UN) and infected Myd88KO iBMDMs. Log_10_ transformed data of p-rS6p mean fluorescent intensity (MFI) is shown where each circle represents data from a single cell. The number for cells analyzed for each condition is noted and the mean for each condition is indicated with a red bar. Statistical analysis was performed with multiple comparisons Kruskal-Wallis one-way ANOVA test. The respective p- values across the different comparisons are shown. **(B-D)** One representative of three biological replicates is shown.

### BinA promotes expansion of the *Legionella*-occupied organelle in an mTOR- dependent manner

One consequence of mTOR induction by *Legionella* is the stimulation of host lipogenesis which satisfies the need for membrane phospholipids as the LCV expands and the bacteria replicate within it. As a result, lipogenesis prevents premature rupture of the intracellular niche and extends the LCV lifespan [4, 5]. Thus, we investigated whether *binA* deletion impacts the mTOR-dependent LCV homeostasis processes. To this end, we first generated a MTOR^−/−^ U937 human monocytic cell line using CRISPR/Cas methodology, which can be terminally differentiated into macrophages following phorbol ester treatment [57] (Fig 3A). To validate the loss of mTOR signaling in the knockout cell line, monocytes were stimulated with *E.coli* LPS (a canonical TORC1 inducer) and the amount of p-rS6p was measured via immunoblot analysis (Fig 3B). In this setting, LPS triggered rS6p phosphorylation in the parental U937 cells but not in the MTOR^−/−^ cells, which is consistent with a loss of mTORC1 activity in the knockout cell line (Fig 3B).

**Fig 3.**
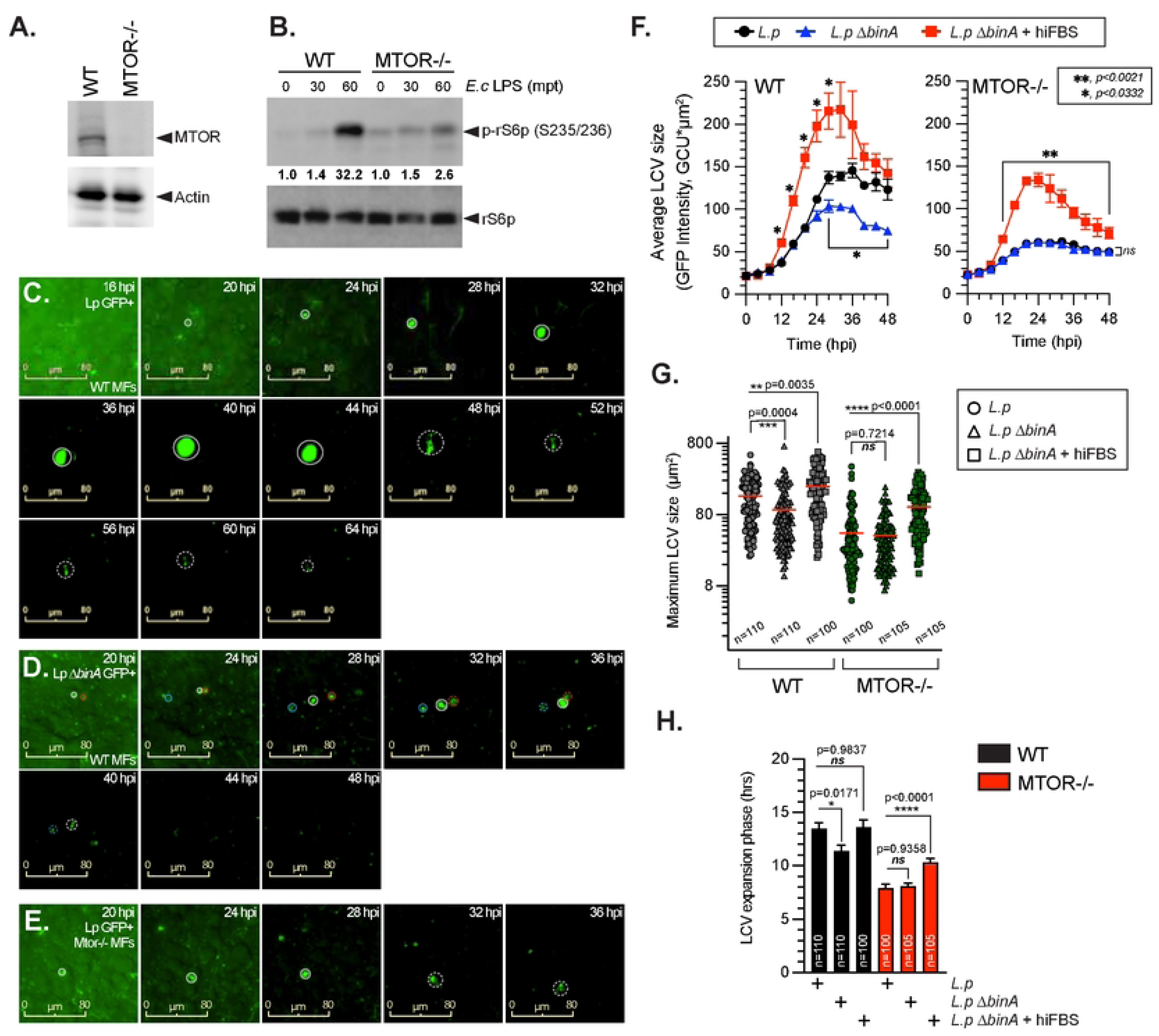
BinA promotes LCV expansion in a MTOR-dependent manner. **(A)** Immunoblot analysis of MTOR abundance in U937 *Mtor* ^−/−^ monocytes and its respective parental WT cell line. **(B)** Loss of TORC1 signaling in U937 *Mtor* ^−/−^ monocytes stimulated with *E.coli* LPS (100 ng/ml) as indicated. **(C-E)** Representative micrograph time-series from live-cell imaging of different GFP+ Lp strains showing LCV dynamics of vacuoles containing either Lp01 **(C)** or Lp01 Δ*binA* **(D)** within U937 macrophages (MFs) or vacuoles occupied by Lp01 inside U937 *Mtor* ^−/−^ MFs **(E)** (MOI = 2.5). Individual LCVs are circled by a solid line prior to and by a dashed line after rupture. Multiple LCVs in the same micrograph are indicated by different line color. **(F)** Population level analysis of LCV size during infection of U937 MFs that express or lack MTOR. The changes in the average LCV sizes ± StDev from three technical replicates are shown. Some conditions were supplemented with hiFBS (5% v/v) to complement the loss of MTOR-dependent metabolite production by the host cells. **(G-H)** Quantitative analysis of LCV dynamics at the level of single LCVs. For each condition, the parameters of at least 100 individually tracked LCVs were analyzed, where each data point is derived from a single vacuole. **(G)** The maximum size reached by individual LCVs under the indicated infection conditions are shown, where the bar indicates population mean value. **(H)** The graph shows the average ± SEM duration of the expansion phase of the pathogen-occupied vacuole for each infection condition. **(F, G-H)** Statistical analysis was completed via two-way **(F)** or one-way **(G-H)** ANOVA with Dunnett’s multiple comparison test using the Lp infection as baseline control. The respective p-values are indicated for each panel. **(B-H)** Data is from one representative of three biological replicates.

Next, U937 macrophages expressing or lacking mTOR were infected with different GFP+ *Legionella* strains and the lifecycle parameters of multiple LCVs from infected macrophages were measured via live-cell imaging (Fig 3C-E). In these infection assays, cells were imaged every 4hrs over several days, which allowed for determination of changes in the LCV size over time as well as the LCV lifespan. LCVs from U937 macrophages infected with *L.p* Δ*binA* GFP+ are shown in Fig 3D, whereas *L.p* GFP+ infections of MTOR+ or MTOR– macrophages are shown in Fig 3C and 3E respectively. In mTOR sufficient macrophages, the LCV expansion rate of vacuoles harboring *binA*− or *binA*+ were identical as demonstrated by the slopes of the LCV growth curves (Fig 3F, left panel). However, the growth of LCVs containing *binA*− bacteria plateaued at a vacuole size that is significantly smaller as compared to LCVs harboring *binA+* bacteria indicative of premature organelle growth arrest (Fig 3F, left panel). Conversely, LCVs harboring *binA*− or *binA*+ bacteria terminated expansion at similar sizes in mTOR deficient macrophages (Fig 3F, right panel) indicating that the BinA-dependent regulation of LCV expansion requires the mTOR kinase. Supplementation of exogenous serum-derived lipids during infection has been shown to rescue the LCV biogenesis defects caused by the loss of mTOR-dependent *de novo* lipogenesis by directly providing lipids for membrane synthesis [4, 5]. Similarly, the expansion of LCVs harboring *binA*− bacteria in both mTOR-sufficient and mTOR-deficient macrophages was restored upon hiFBS supplementation further supporting the notion of phenotypic link between the absence of BinA and mTOR activities (Fig 3F). Single vacuole analysis data showed that on average the maximum size achieved by individual bacteria-occupied organelles was significantly lower when either MTOR or *binA* were knocked out (Fig 3G) and organelle expansion terminated earlier as well (Fig 3H). Importantly, both of these LCV biogenesis parameters were indistinguishable between infections with *binA*− and *binA*+ bacteria in the absence of mTOR activity (Fig 3G-H).

Taken together these data indicate a mechanistic connection between BinA and mTOR signaling that phenotypically manifests specifically in premature termination of LCV expansion and an overall lower bacterial load per organelle that can be rescued by exogenous serum-derived lipids supplementation.

### BinA sensitizes mTOR signaling in response to amino acid stimulation upstream of the heterodimeric Rag GTPase complex

To gain insight into the mechanism by which BinA regulates mTOR, we first tested whether BinA is sufficient to trigger TORC1 activation upon amino acids stimulation when produced by eukaryotic cells independent of infection. To this end, HK293 cells ectopically producing GFP or GFP-tagged BinA were assayed for their capacity to trigger TORC1 signaling in a prototypical amino-acid withdrawal/re-stimulation assay where cells were cultured in amino acid-free media for 4 hours and subsequently were treated for 30min with various amounts of arginine (Fig 4A) or leucine (Fig 4B). As expected, treatment with either amino acid triggered phosphorylation of S6K (a TORC1 substrate) and its immediate downstream substrate rS6p (Fig 4A-B) indicative of TORC1 activation. Notably, expression of GFP-BinA but not GFP resulted in TORC1 activation at lower concentrations of Arg (10 - 50µM vs. 1000µM) (Fig 4A) and Leu (10 – 100µM vs. 200 – 400µM) (Fig 4B). In addition, BinA expression alone did not trigger phosphorylation of either S6K or rS6p (Fig 4A-B) in the absence of Arg/Leu indicating that the TORC1 pathway retains stimulus-dependency despite the BinA-dependent sensitization. Furthermore, kinetic analysis of TORC1 activation by different doses of Arg in the presence of BinA revealed increased p- rS6p signal at early time points following stimulation, which is consistent with an accelerated and a sensitized response (Fig 5A). Similar results were observed following Leu stimulation (Fig 5B). Based on these data we conclude that BinA directly or indirectly lowers the threshold for TORC1 activation upon stimulation with either Arg or Leu in the absence of other bacterial factors.

**Fig 4.**
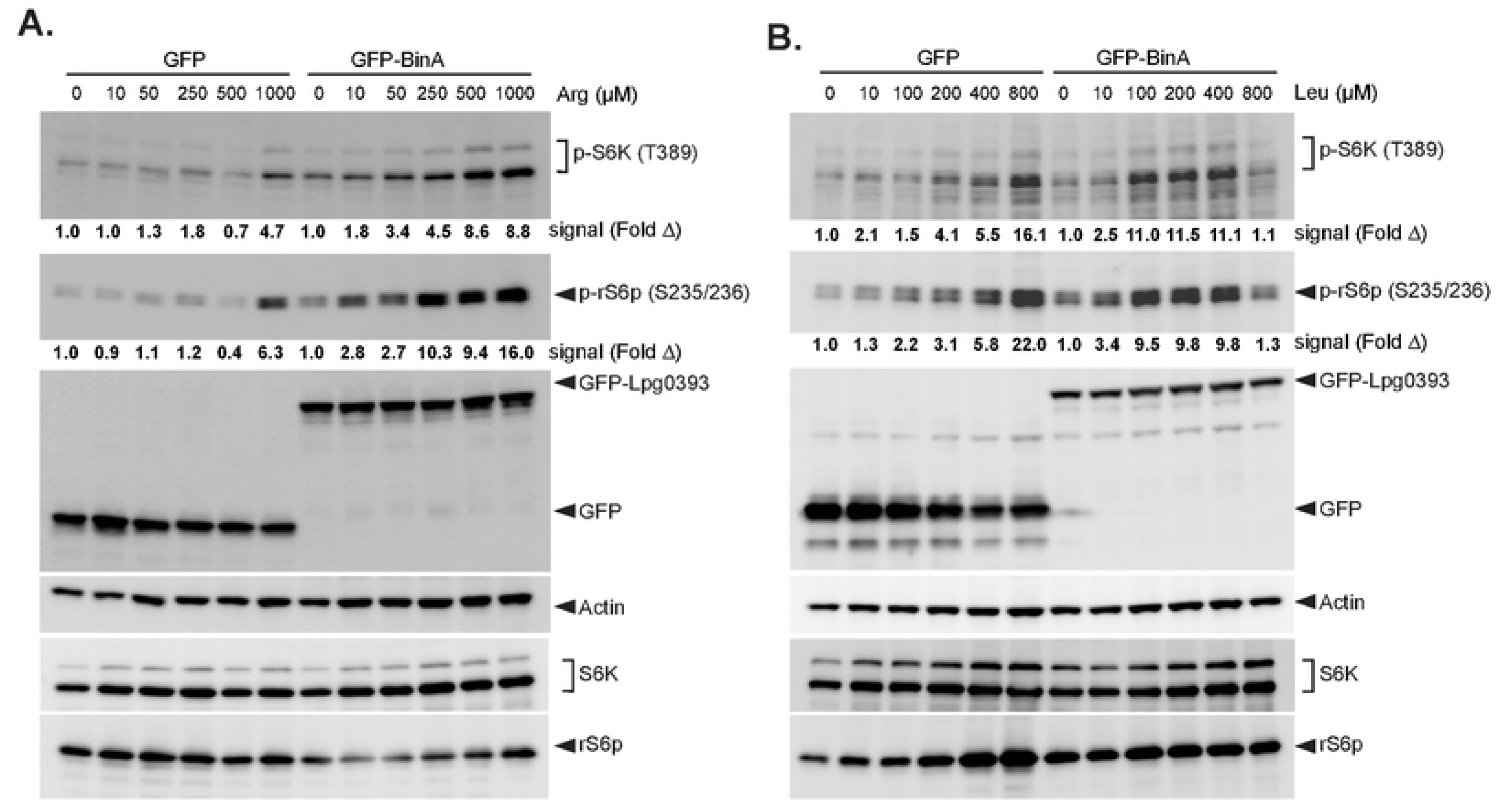
BinA sensitizes TORC1 signaling downstream of arginine and leucine stimulation. **(A-B)** Ectopic expression of BinA in eukaryotic cells is sufficient to sensitize TORC1 signaling induced by arginine or leucine. Data shows dose-dependent TORC1 activation by Arg **(A)** or Leu **(B)** at 30 min post-treatment (mpt) in amino-acid starvation/refeeding experiments using HK293 cells transfected with either GFP or GFP- BinA. **(A-E)** Band signal intensity in the phospho-immunoblots for S6K (A-B) and rS6p (A- E) for each condition was quantified and is presented below each figure panel as fold change from untreated cells. The data shown is from one experiment out of at least 3 biological replicates.

**Fig 5.**
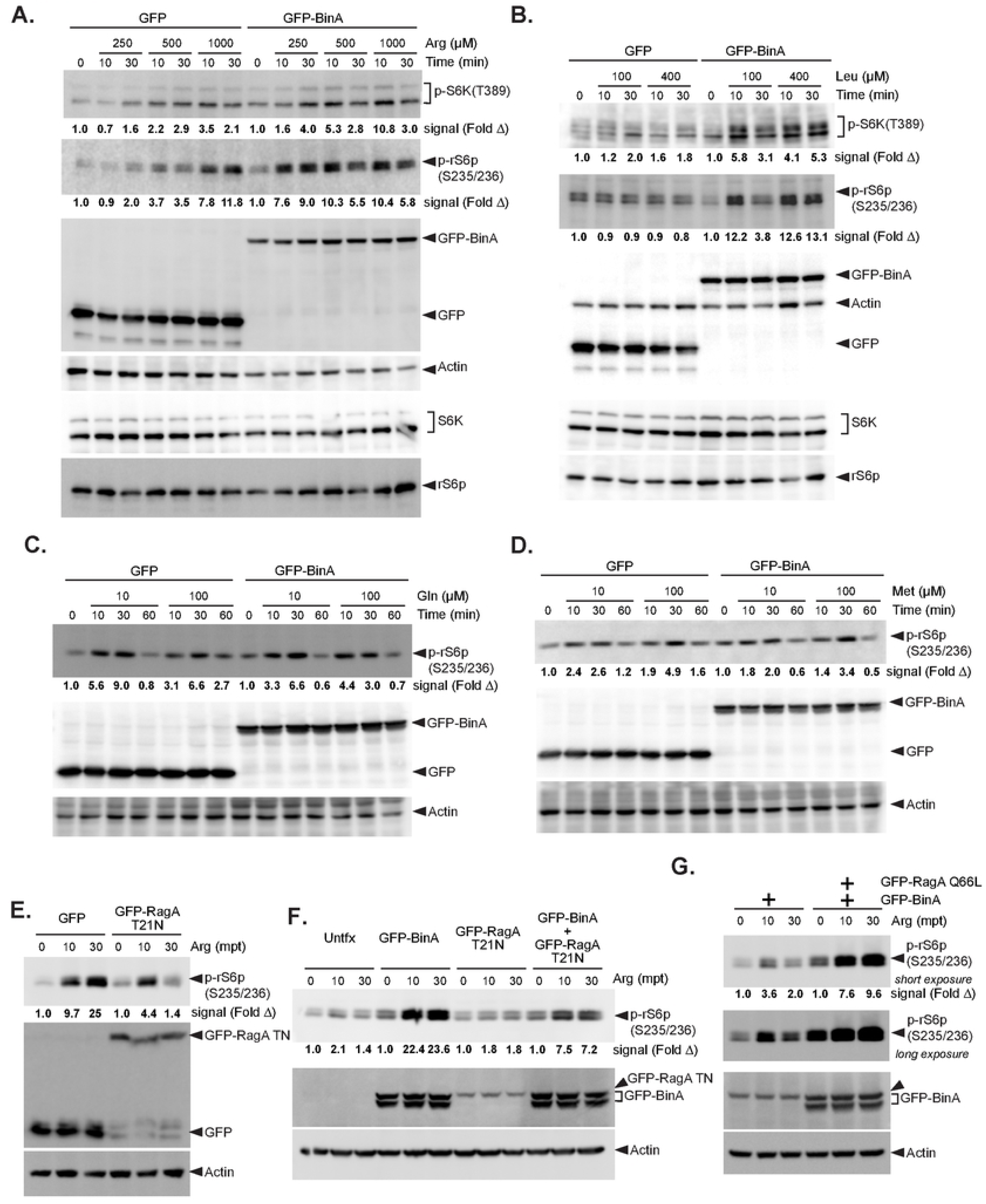
Sensitization of TORC1 signaling by BinA is stimulus-dependent and requires the Rag GTPase complex. Comparison of TORC1 signaling kinetics in response to stimulation with different doses of arginine **(A)**, leucine **(B)**, glutamine **(C)** or methionine **(D)** in HK293 cells that ectopically produce either GFP or GFP-BinA. **(E-F)** Ectopic expression of the GDP-locked RagA T21N allele inhibits arginine-induced TORC1 signaling in HK293 cells lacking **(E)** or producing GFP-BinA **(F)** in amino-acid starvation/refeeding experiments. Treatments with either 1mM or 250 µM Arg are shown respectively in **(E)** and **(F)**. **(G)** The GTP-locked RagA Q66L allele augments TORC1 signaling in BinA-expressing HK293 treated with low-dose 100 µM Arg for the indicated time periods in a starvation/refeeding experiment. **(A-G)** Immunoblot data from amino acid starvation/refeeding experiments are shown, where band signal intensity in the phospho- immunoblots for S6K **(A-B)** and rS6p **(A-G)** for each condition was quantified and is presented below each figure panel as fold change from untreated cells. The data shown is from one experiment out of three biological replicates.

The regulatory network coordinating TORC1 signaling involves multiple amino acid-specific transporters, sensors and signaling auxiliary intermediates [20, 58]. We exploited the inherent network complexity to begin addressing the BinA-driven TORC1 sensitization mechanism. First, we tested whether BinA function is broadly prevalent in multiple TORC1-activation pathways, which would likely suggest direct regulation of the TORC1 complex. To this end, we took advantage of two TORC1 amino acid agonists – Met and Gln - that are sensed via mechanisms distinct from Leu and Arg [58–61]. Broadly speaking, sufficiency of different amino acid is detected and relayed to the TORC1 complex via multiple pathways that incorporate distinct small GTPases (RagA/B/C/D, Rab1a or Arf1) which function as molecular switches to recruit TORC1 to either lysosomal or Golgi membranes for activation [20, 21, 58, 60, 62, 63]. S-adenosylmethionine (SAM) – a methionine-derived metabolite – stimulates TORC1 in a Rag GTPase complex-dependent manner similarly to Arg and Leu [59]. Conversely, glutamine stimulates TORC1 through an Arf1-dependent mechanism [60]; however, glutamine breakdown via glutaminolysis also produces α-ketoglutarate, which can stimulate TORC1 in Rag GTPase complex-dependent manner [61]. Thus, we reasoned that TORC1 stimulation by Met and Gln in the context of BinA could provide mechanistic insight into the sensitization process. To this end, HK293 cells expressing GFP or GFP-BinA were stimulated with different amounts of Gln (Fig 5C) or Met (Fig 5D). Unlike Leu and Arg stimulation, TORC1 signaling was not sensitized by BinA after treatment with either Gln or Met (Fig 5C-D) indicating that BinA unlikely targets the TORC1 complex directly which would result in sensitization to all amino acid treatments. These data also indicate that BinA likely sensitizes the Rag GTPase complex-dependent TORC1 activation pathway, idea that was tested in the next set of experiments.

The heterodimeric Rag GTPase complex coordinates TORC1 signaling by recruiting TORC1 to the lysosomal membrane based on amino acids availability where the complex is directly activated by the small GTPase Rheb [20, 58]. The primary components of the Rag complex are RagA, RagB, RagC and RagD which form a heterodimer of a small subunit (RagA or RagB) and a large subunit (RagC or RagD) [21, 63]. Under amino acid replete conditions, the complex consists of a GTP-bound RagA/B and a GDP-bound RagC/D, whereas upon amino acid starvation the nucleotide-bound state of the GTPases is reversed [20, 58]. Ectopic expression of different point mutant RagA/B/C/D alleles that lock the GTPase in distinct nucleotide bounds states promotes or interferes with TORC1 signaling depending on the allele [21, 63]. As expected, expression of the GDP-locked point mutant allele RagA T21N interfered with Arg-induced TORC1 activation (Fig 5E). Cells co-expressing RagA T21N and BinA exhibited ∼ 3-fold reduction in the p-rS6p signal in low dose Arg starvation/refeeding experiments as compared to cells expressing BinA alone (Fig 5F) indicating that TORC1 sensitization by BinA is a Rag GTPase complex-dependent event. Consistent with that idea, cells producing BinA together with the GTP-locked RagA Q66L allele hyperactivated TORC1 when stimulated with low dose Arg in comparison to cells producing RagA Q66L alone (Fig 5G).

Based on these data, we conclude that (i) BinA sensitizes TORC1 signaling specifically to Arg/Leu stimulation by targeting the major Rag GTPase complex-dependent arm of the pathway at a point upstream of the TORC1 complex and (ii) BinA can enhance TORC1 signaling independent of other bacterial factors.

### BinA enhances amino acid uptake in infected macrophages

We reasoned that BinA likely regulates a factor or a process common to the pathways relaying arginine and leucine sufficiency to TORC1 because response sensitization was specific for Arg and Leu stimulation but not Met or Gln treatment (Fig 5A-D). The methionine-derived metabolite SAM triggers TORC1 activation through the sensory proteins SAMTOR (S-adenosylmethionine sensor upstream of mTORC1) and the Rag GTPase complex [59], thus it is unlikely BinA targets the following proximal upstream TORC1 regulators that are shared by the Arg, Leu and Met sufficiency response pathways – (i) the pentameric Ragulator complex that tethers the Rag GTPase complex to lysosomes [23]; (ii) the GATOR complex (GTPase activating proteins toward Rags), which has GAP activity towards the small RagA/B GTPases [64]; (iii) the Rag GTPase complex, which recruits TORC1 to lysosomes [21, 63]. Therefore, we hypothesized that BinA might regulate Leu/Arg transporters or the respective amino acids cytosolic sensors.

TORC1 activation through the amino acid sufficiency pathway can be triggered independently of amino acid transport pathways by the lysosomotropic agent LLoMe (l-leucyl-l-leucine methyl ester). LLOMe is internalized via receptor mediated endocytosis and in lysosomes is converted into a hydrophobic polymer which subsequently transverses the lysosomal membrane and triggers TORC1 signaling via the amino acid sensing pathway [65]. BinA expression did not increase nor accelerate TORC1 signaling in HK293 cells upon stimulation with different doses of LLoMe (Fig 6), which would be expected if BinA functions downstream of amino acid import. Because BinA cannot sensitize TORC1 when amino acid transport is eliminated as a variable, the data suggests BinA may function by regulating Arg/Leu import.

**Fig 6.**
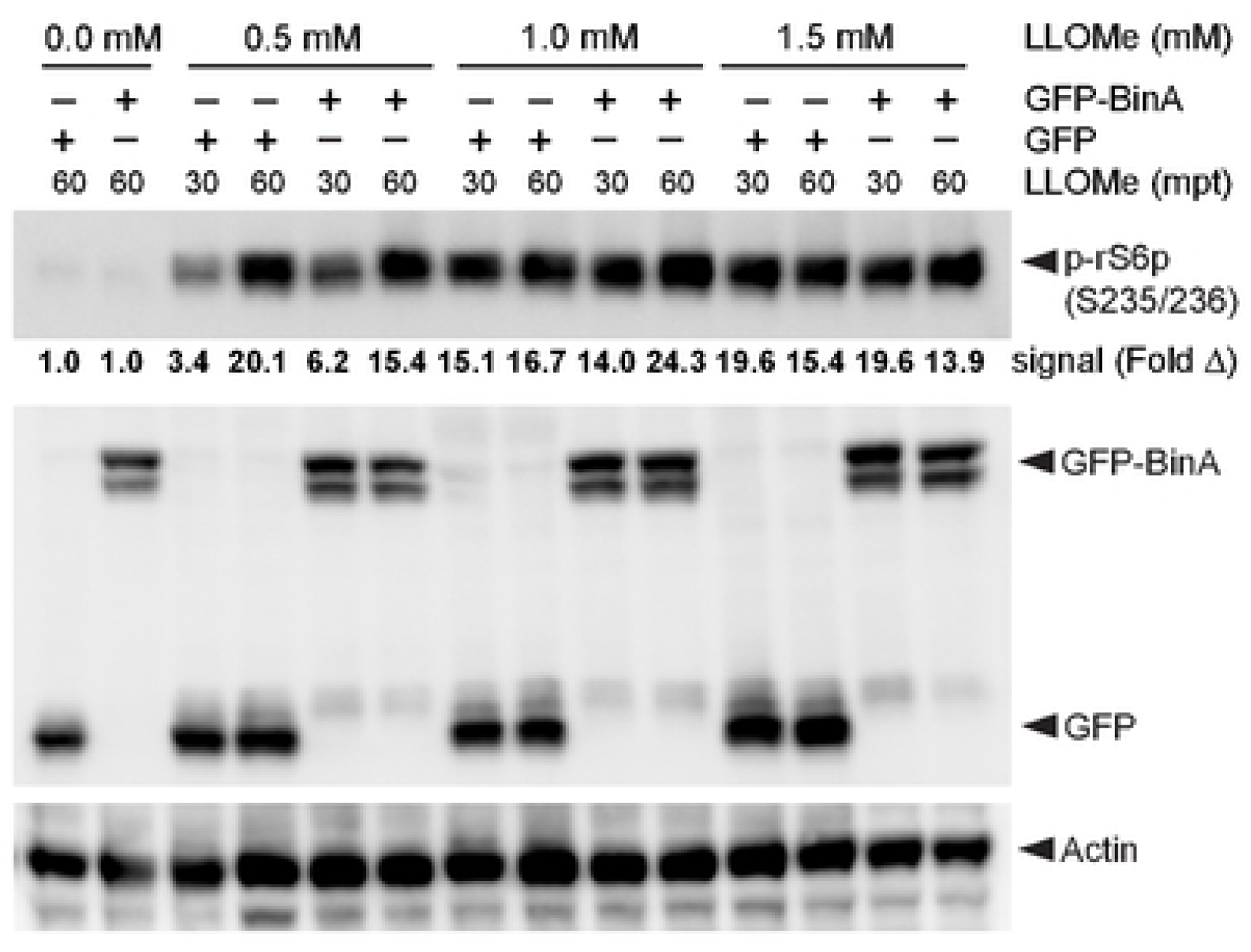
BinA does not sensitize TORC1 signaling in response to L-Leucyl-L-Leucine methyl ester. MTOR sensitization by BinA is not evident when HK293 cells expressing either GFP or GFP-BinA are treated with low doses of the membrane permeant TORC1 signaling agonist L-Leucyl-L-Leucine methyl ester (LLOMe). Immunoblot data from amino acid starvation/LLOMe refeeding experiment are shown, where band signal intensity in the phospho-immunoblot for rS6p for each condition was quantified and is presented below the respective figure panel as fold change from untreated cells. The data shown is from one experiment out of three biological replicates.

Therefore, BinA capacity to modulate amino acid import was investigated in the context of infection. To this end, we utilized a fluorescent sensor to measure specifically the uptake of the amino acid analogue L-BPA (L-boronophenylalanine) which is imported by the heterodimeric branched amino acid transporter LAT1 (L-type amino acid transporter 1) (Fig 7A) [66]. For these experiments, Myd88KO immortalized BMDMs (iBMDMs) were infected with *binA*+ or *binA*− strains and L-BPA import by infected cells was measured for 5min at 4 hpi (Fig 7B). Macrophages infected by the *L.p* Δ*binA* imported significantly lower amounts of L-BPA as compared to the parental wild type strain (Fig 7B). Importantly, plasmid-borne BinA complemented the L-BPA import defect demonstrating that BinA secreted by *L.p* during infection can alters amino acid transport through LAT-1 (Fig 7B). As expected, treatment with the LAT-1 inhibitor BCH (2-aminobicyclo-(2,2,1)-heptane-2- carboxylic acid) reduced L-BPA import (Fig7B). Together, these data implicate BinA as a bacterial regulator used by *L.p* to enhance amino acid import in the cytosol of infected cells which can account for BinA capacity to sensitize TORC1 signaling.

**Fig 7.**
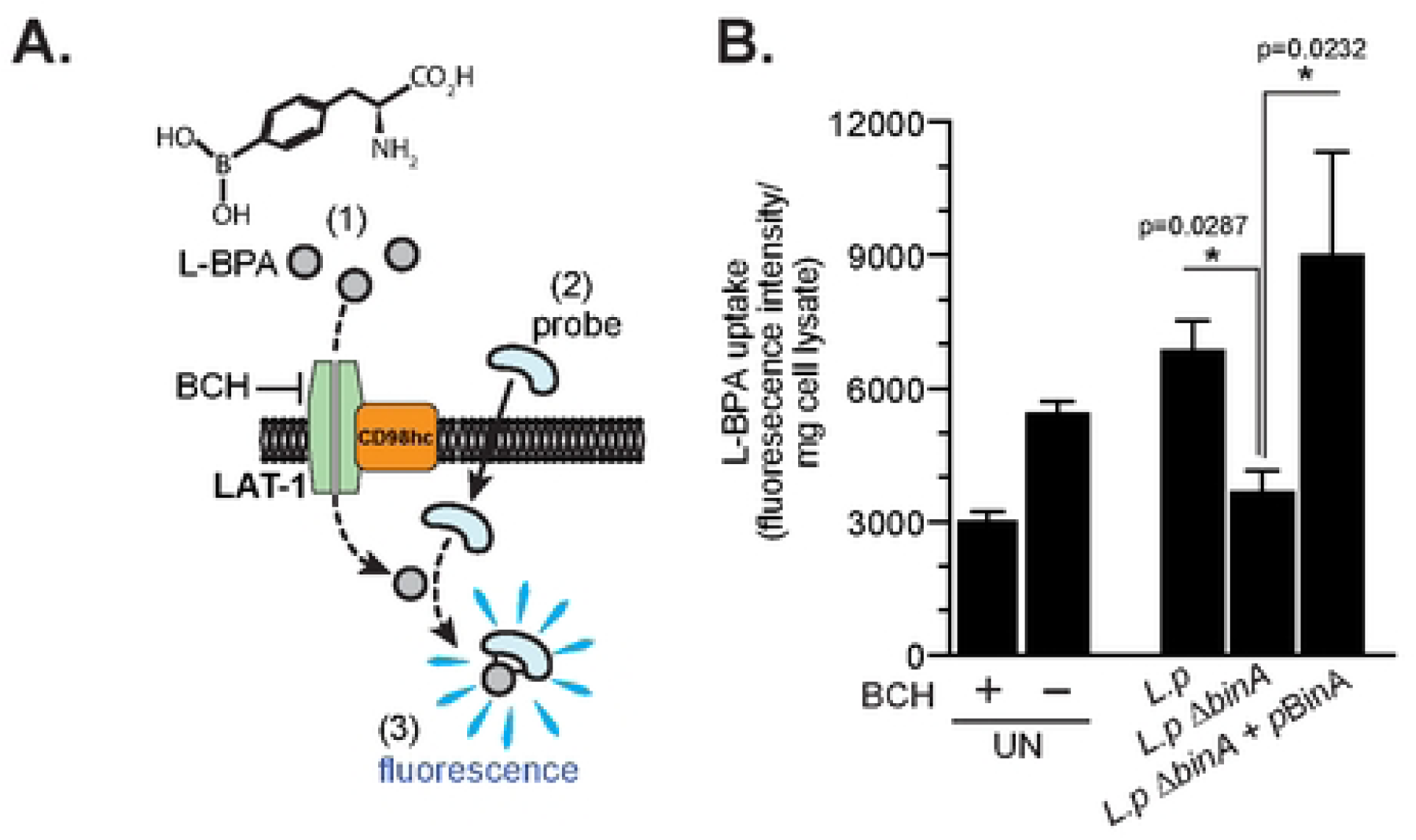
BinA is required for optimal amino acid import through the L-type amino acid transporter 1 (LAT1) in macrophages infected with Lp. **(A)** Schematic depiction of the LAT1-dependent uptake assay for import of the amino acid analog ρ-Boronophenylalanine (L-BPA). Cell were treated with L-BPA for 5min (1) after which cells were extensively washed, followed by the addition of a specific membrane permeable probe for additional 5 min (2), which emits fluorescent signal when bound to L-BPA(3). **(B)** Graph shows L- BPA uptake by Myd88 KO iBMDMs infected with the indicated Lp strains at MOI=20 or left uninfected (UN). In some control conditions, the LAT1 inhibitor 2-Aminobicyclo [2.2.1] heptane-2-carboxylic acid (BCH) at 1mM was added 10 min prior to and for the duration of the L-BPA uptake assay. BPA uptake lasted 5min and was measured at 4 hpi, after which fluorescence output was measured, the cells were lysed and total protein content was measured. Mean fluorescence data was normalized to the total protein content. Averages ± SEM of data from three biological repeats are shown for each condition. Statistical analysis was completed via one-way ANOVA Kruskal-Wallis test with Dunn’s multiple comparison and the respective p-values among the conditions compared are indicated.

Previous data indicates that *Legionella*-induced inhibition of host protein synthesis triggers TORC1 signaling in infected cells by increasing the cytosolic amino acid pool [48]; however, it is unclear if import of extracellular amino acids pool contributes to TORC1 activation by *L.p*. Thus, we measured TORC1 signaling under infection conditions in the presence or absence of extracellular amino acids. To this end, Myd88KO iBMDMs were infected for 4hrs in medium containing amino acids (serum free RPMI), which resulted in robust 15.3-fold induction of p-rS6p as compared to uninfected cells (Fig 8A). Subsequently, infections were continued either in the presence (serum free RPMI) or absence (HBSS) of extracellular amino acids for additional 4 hours (Fig 8A). In this setting TORC1 signaling gradually decreased upon withdrawal of extracellular amino acids reaching the level observed in uninfected cells within 2 hours of starvation demonstrating that the non-cytosolic amino acid pool is needed from sustained TORC1 activation in *L.p* infections (Fig 8A). In agreement, TORC1 signaling was maintained in the presence of exogenous amino acids (Fig 8A). The phosphorylation of eIF2α (Eukaryotic initiation factor-2α), which is triggered by cellular stress responses, did not decline under the same conditions indicating that amino acid withdrawal did not reduce macrophage capacity to carry out signal transduction in general (Fig 8A).

**Fig 8.**
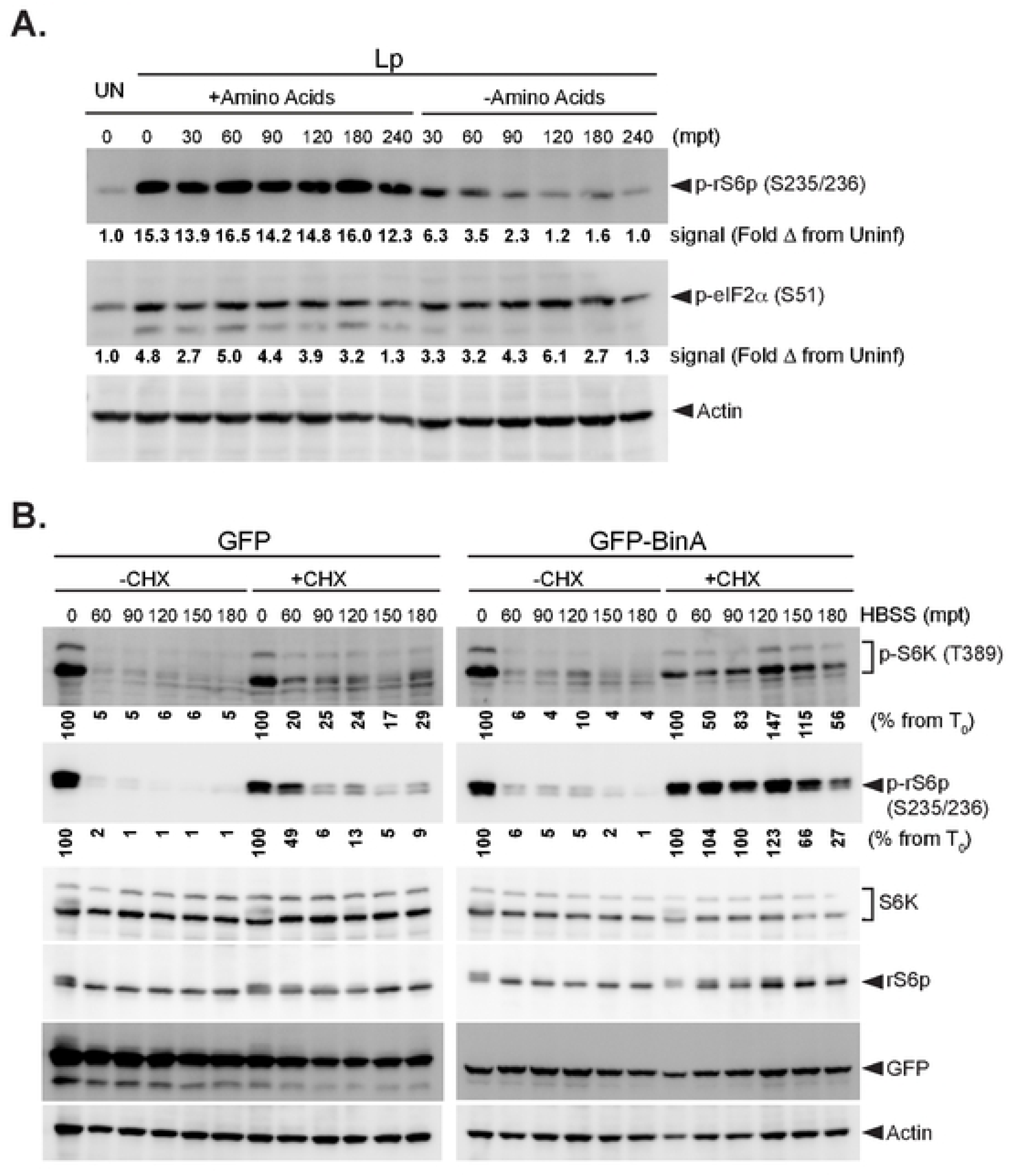
Extracellular amino acids sustain TORC1 activity in Lp infected cells. **(A)** Immunoblot analysis of TORC1 (p-rS6p) and stress response (p-eIF2α) signaling in Lp infected macrophages in the presence or absence of extracellular amino acids (MOI=15). Data is derived from Myd88 KO iBMDMs that infected with Lp for 4 hours after which RPMI-derived amino acids were either kept or withdrawn by replacing the RPMI medium with amino-acid free Hank’s Balanced Salt Solution (HBSS) for the indicated time periods. Band signal intensity for p-rS6p and p-eIF2α for each condition was quantified and is presented below the respective figure panels as fold change from uninfected (UN) cells. **(B)** BinA expression prolongs TORC1 activation caused by protein translation inhibitors. Changes in TORC1 signaling upon amino-acid withdrawal (HBSS treatment) from HK293 cells producing either GFP or GFP-BinA in the presence or absence of 350µM cycloheximide (CHX). CHX was added 30min prior to and was retained during the HBSS treatment. Band signal intensity for p-S6K and p-rS6p for each condition was quantified and is presented below the respective figure panels as percentage of signal from cells that were not treated with HBSS. The data shown is from one experiment out of three biological replicates.

Next, we investigated the interplay between the cytosolic and the extracellular amino acid pool in TORC1 signaling in the context of BinA expression. Treatment of HK293 cells expressing either GFP or GFP-BinA with the protein translation inhibitor cycloheximide, which increases the cytosolic amino acid pool [51], prolonged TORC1 signaling upon withdrawal of extracellular amino acids (Fig 8B). However, under those conditions the p-rS6p signal in cell lacking BinA was reduced by >90% within 90 mpt with HBSS, whereas in cell expressing BinA the p-rS6p signal started declining only after 150 mpt (Fig 8B). Conversely, TORC1 signaling was rapidly shut off when extracellular amino acids were withdrawn in the absence of cycloheximide treatment regardless of BinA expression (Fig 8B). Taken together, these data indicate BinA can prolong TORC1 signaling upon withdrawal of extracellular amino acids providing the cytosolic amino acid pool is not depleted by the protein translational machinery. Because (i) amino acid transporters localize at the plasma membrane as well as along the endolysosomal continuum [67–69] and (ii) amino acid transport is in part linked through antiporter activities which couples import of one amino acid with the export of another [70, 71], the BinA- enhanced import of TORC1 stimulating amino acids, such as Leu and Arg, might depend on increased cytosolic amino acid pool for sustained bi-directional amino acid transport.

### BinA sensitizes TORC1 signaling by altering endosomal trafficking determinants

BinA has been shown *in vitro* to activate the Rab5 GTPase family members – Rab5, Rab21 and Rab22 – by catalyzing nucleotide exchange via its VPS9-like domain [56]. Rab5 GTPase family members specialize in the regulation of multiple membrane trafficking events along the endocytic compartment [72]. Rab22 intercepts and rapidly redirects endocytosed CIE (Clathrin independent endocytosis) cargo – including CD98hc- containing heterodimeric transporters – to recycling endosomes, whereas Rab21 controls recycling of CIE cargo from recycling endosomes to the cell surface [73–76]; conversely, Rab5 promotes endosomal maturation [72]. Genetic evidence indicates that TORC1 responses to amino acids sufficiency requires Rab5 presumably because of endocytic maturation [77]. Given the role of Rab5 GTPase family members in trafficking of amino acid transporters and BinA capacity to promote amino acid import to sensitize TORC1, we investigated whether Rab5, Rab21 or Rab22 are involved. To this end, TORC1 signaling was assessed after arginine stimulation in HK293 cells expressing BinA alone or in combination with different Rab5/21/22 alleles, including constitutively active GTP-locked as well as dominant negative GDP-locked mutants. For Rab5b, neither the GTP-locked Q79L nor the GDP-locked S34N mutant impacted BinA capacity for TORC1 sensitization in response to Arg and Leu (Fig 9A-B and Fig S1A-B). Conversely, expression of either the GDP-locked Rab21 T31N or Rab22a S19N inhibited TORC1 sensitization by BinA in response to Arg (Fig 9C-F) and Leu (Fig 9D and 9F and Fig S1C-F). Based on these finding, we conclude that Rab21 and Rab22 but not Rab5 mediate TORC1 sensitization by BinA after Arg or Leu stimulation. Both Rab21 and Rab22 dominant negative alleles independently ablated TORC1 signaling indicating that these GTPase integrate linearly within the same regulatory pathway controlled by BinA. Because Rab21 and Rab22 function in endocytic cargo transport to and from the recycling endosomal compartment, including amino acid transporters, it is likely that BinA-dependent mTOR sensitization activity also originates at that location.

**Fig 9.**
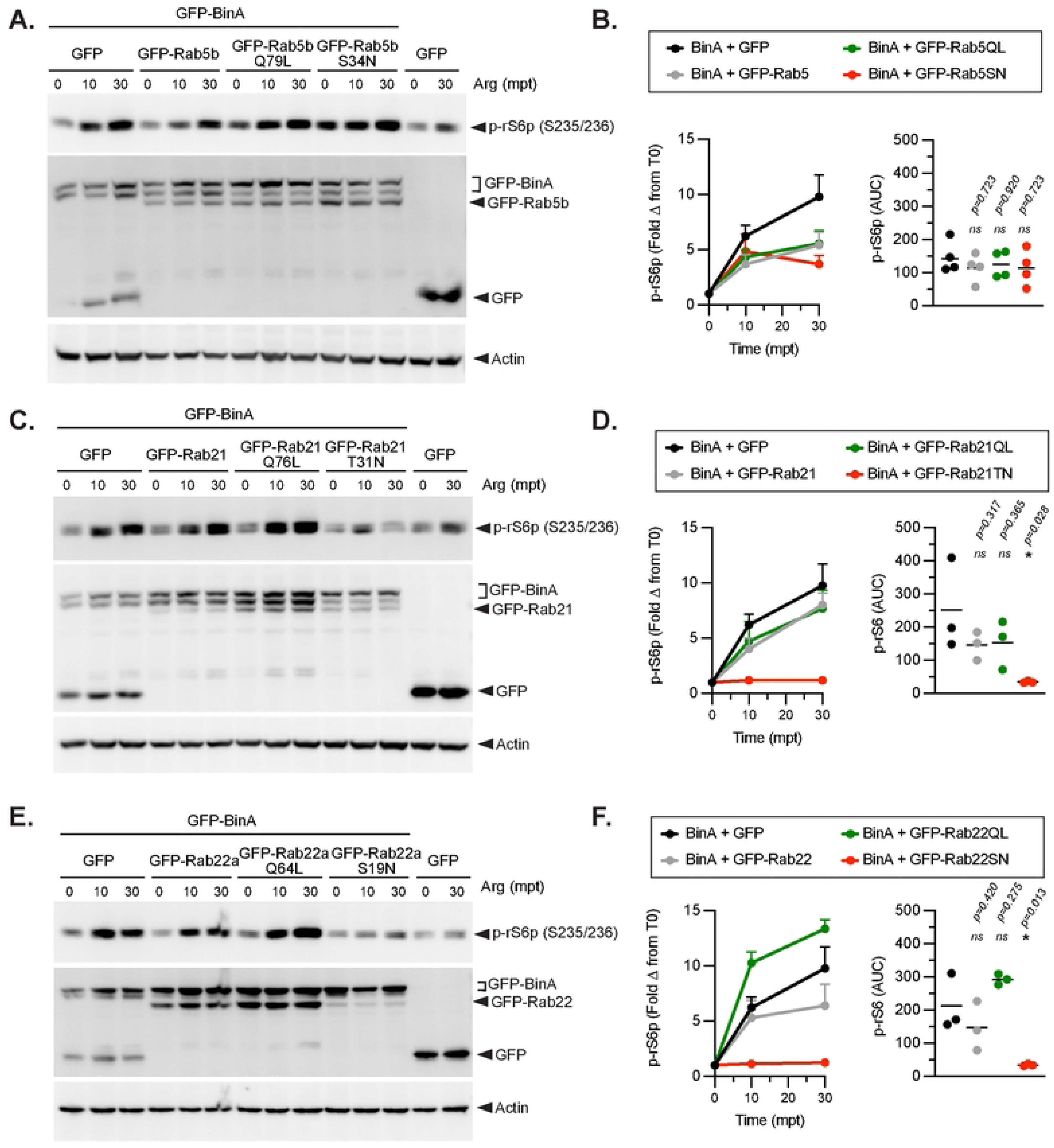
Sensitization of arginine-dependent TORC1 signaling by BinA is regulated by Rab21 and Rab22a but not Rab5. Kinetics of TORC1 activation triggered by starvation/refeeding stimulation with 500µM Arg in HK293 cells producing BinA in the presence or absence of different Rab5 **(A-B)**, Rab21 **(C-D)**, or Rab22a **(E-F)** alleles. **(B, D and F)** Quantitative analyses of band signal intensities show Averages ± StDev from at least three biological repeats (left panels) and the respective area-under-the curve (AUC) analyses are presented in the right panels. Statistical analyses were completed with one- way ANOVA with Dunnett’s multiple comparison test using the ‘BinA+GFP’ as control group and p-values are indicated in the respective data panels.

These data were surprising because previous study on the role of different Rabs in amino acid-induced TORC1 activation uncovered requirement for Rab5 but not Rab21 or Rab22 [77]. To investigate whether BinA dictates the observed mechanistic differences – i.e. the switch from a Rab5 dependency to Rab21/22 dependency – we took advantage of the RagA GTP-locked mutant Q66L as an alternative eukaryotic TORC1 sensitization factor. Co-expression of RagA Q66L with Rab5b S34N significantly reduced TORC1 activation by either Arg (Fig 10A) or Leu (Fig 10B). Conversely, no significant phenotype was observed upon amino acid stimulation in cells expressing RagA Q66L together with either Rab21 T31N or Rab22a S19N (Fig 10A-B). Thus, signaling upstream of TORC1 appears to switch from Rab21/22-dependency to Rab5-dependency contingent upon the presence of BinA or RagA Q66L respectively. Interestingly, when Rab-dependency studies utilizing the Rab5/21/22 GDP-locked mutants were repeated in HK293 cells that did not produce either BinA or RagA G66L, TORC1 activation by Arg (Fig 11A-C) or Leu (Fig S2A-C) was not affected. A functional redundancy in the amino sufficiency detection and response pathways downstream of Arg and Leu can reconcile the data, where parallel Rab5-dependent and Rab21/22-dependent cascades generate amino acid sufficiency signaling input to the Rag GTPare complex.

**Fig 10.**
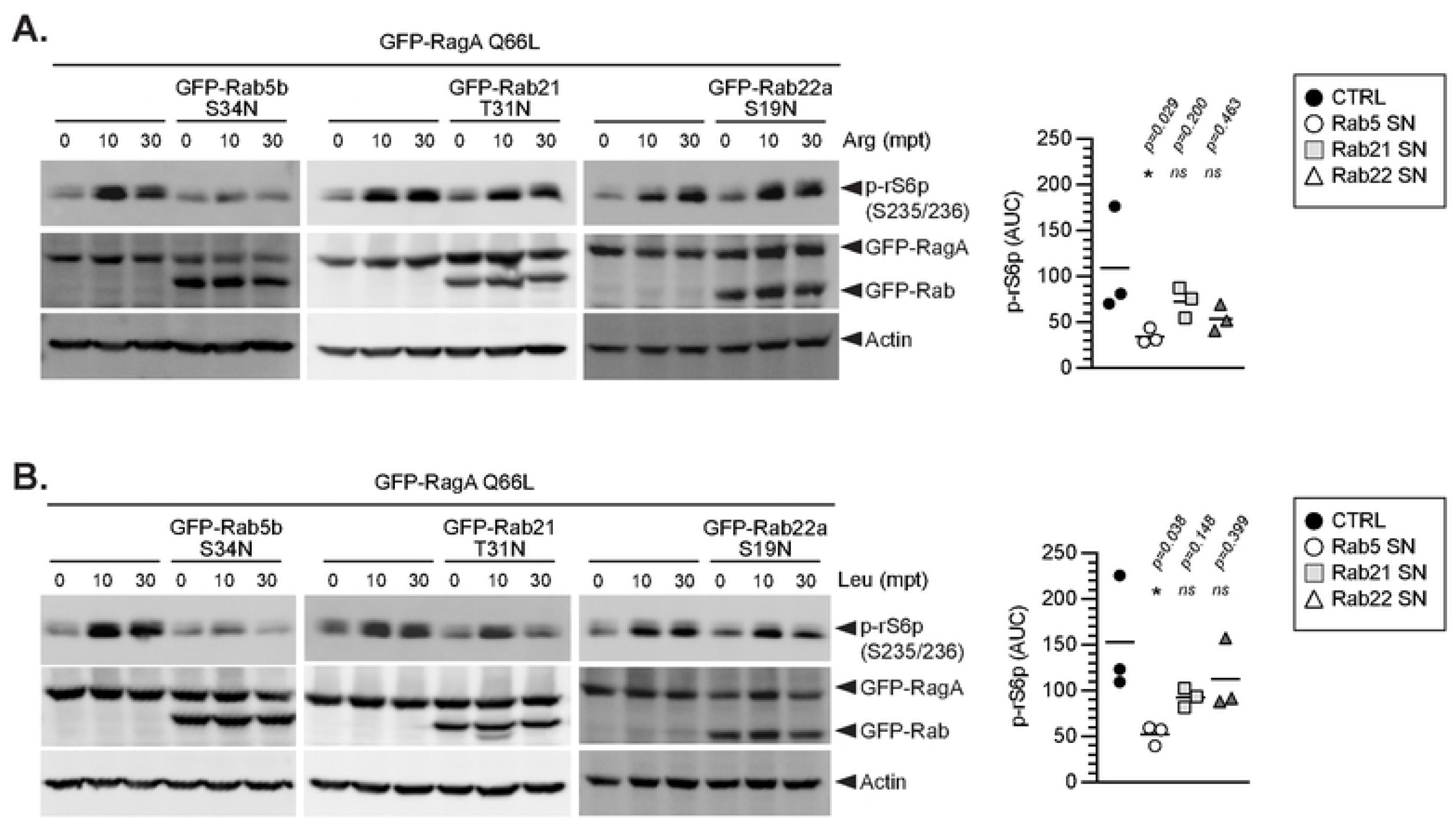
Sensitization of TORC1 signaling by the RagA Q66L allele is regulated by Rab5 but not Rab21 or Rab22a. Kinetics of TORC1 activation triggered by starvation/refeeding stimulation with either 500µM Arg **(A)** or 100 µM Leu **(B)** in HK293 cells producing RagA Q66L in the presence or absence of different GDP-locked Rab5, Rab21 or Rab22a alleles. **(A-B)** Immunoblots from representative experiments are shown in the left panels and AUC analyses from three biological repeats are graphed in the right panels. Statistical analyses were completed with one-way ANOVA with Dunnett’s multiple comparison test using the ‘GFP-RagA Q66L’ as control group and p-values are indicated in the respective data panels.

**Fig 11.**
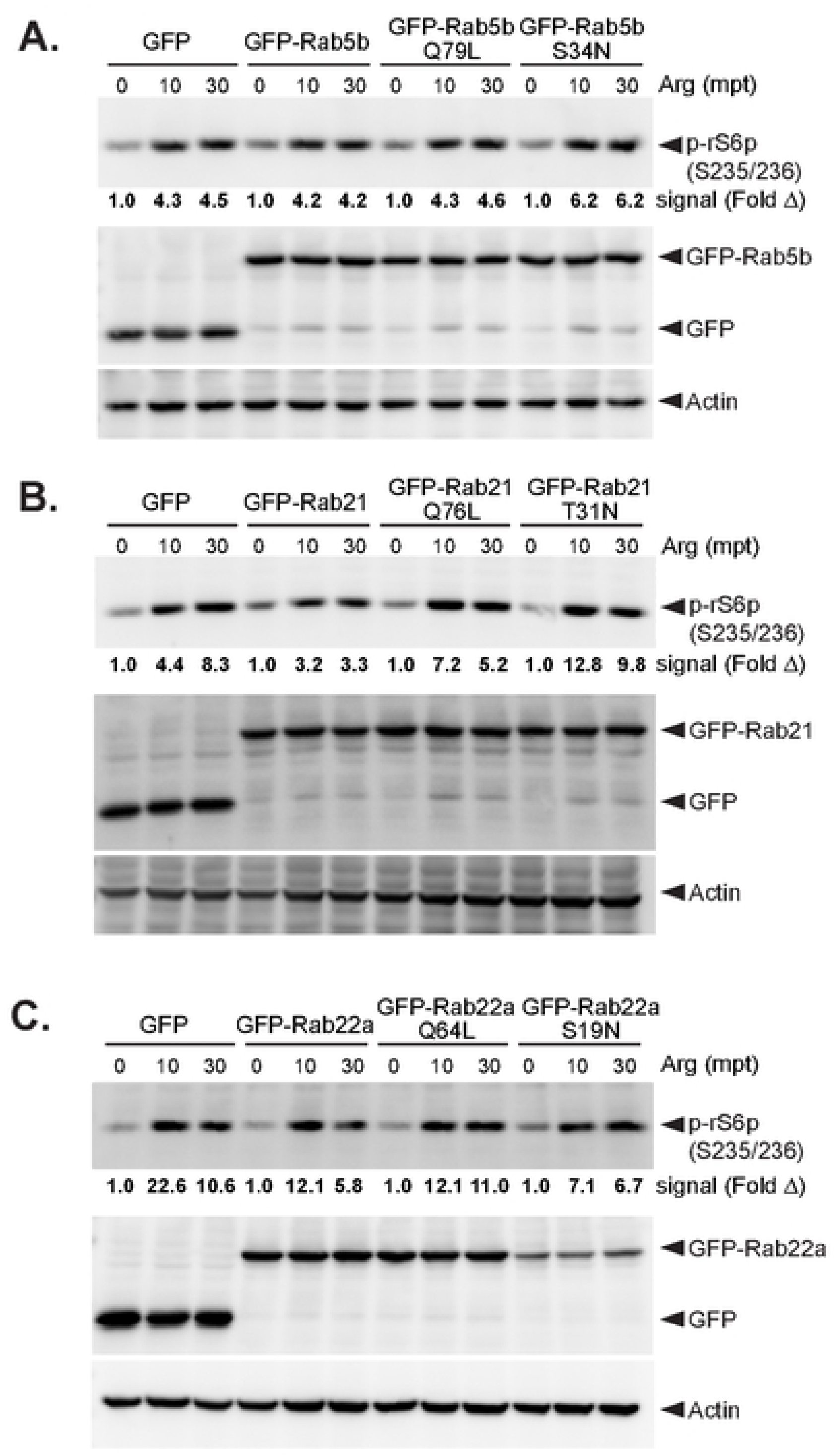
Rab5 family GTPs are dispensable for TORC1 signaling triggered by arginine in the absence of BinA or RagA Q66L sensitization. Kinetics of TORC1 activation triggered by starvation/refeeding stimulation with 1mM Arg in HK293 cells producing GFP or different Rab5 **(A)**, Rab21 **(B)**, or Rab22a **(C)** alleles. Band signal intensity in the phospho-immunoblot for rS6p for each condition was quantified and is presented below the respective figure panel as fold change from untreated cells. The data shown is from one experiment out of three biological replicates.

In amino acid starvation/refeeding experiments which included Rab22 alleles the amount of the Rab22a S19N mutant detected via immunoblotting was consistently lower compared to Rab22a or Rab22a Q64L (Figs 9E, 11C and Figs S1E, 2C). This phenomenon was not observed when lysates from cells cultured in nutrient rich conditions were prepared (data not shown). Directly comparison between Rab22a S19N amounts in HK293 cells prior to and 4 hours post amino acid withdrawal confirmed a reduction by more than 70% (Fig S3), indicating a potential link between amino acids sufficiency sensors and the Rab22 nucleotide bound state, where inability to cycle from the GDP-bound state under amino acids starvation targets the protein for degradation.

### BinA GEF activity is dispensable for TORC1 sensitization

Given BinA GEF activity towards Rab21/Rab22 [56] and the dependence of its TORC1 sensitization capacity on Rab21 and Rab22 we investigated if BinA GEF activity is required for TORC1 sensitization. The BinA D41A allele contains a loss-of-function mutation in the VPS9 domain at a residue required for the nucleotide exchange activity [56]. Indeed, HK293 cells expressing BinA D41A lacked enlarged Rab5-decorated endosomes unlike cells expressing the wild type BinA allele (Fig S4A-C). Expression of BinA D41A by HK293 cells augmented TORC1 signaling in response to Arg comparably if not slightly better than the wild type allele indicating that the GEF activity of the VPS9 domain is dispensable for TORC1 sensitization (Fig 12A). To determine whether that was the case under infection conditions as well, the *L.p* Δ*binA* mutant strain was complemented with an IPTG-inducible plasmid-borne copy of BinA D41A (Fig 2A). In macrophage infections, *binA* deletion strains complemented with either the GEF-sufficient or the GEF-deficient D41A allele restored TORC1 signaling as indicated by the robust p- rS6p signal measured by quantitative microscopy (Fig 12B-D). In those assays, higher p- rS6p signal was detected on average in cells harboring bacteria expressing the D41A mutation as compared to the parental allele even though protein expression was similar (Fig 2A). Based on our data, we conclude that TORC1 sensitization by BinA is independent of the GEF activity of the VPS9 domain towards the Rab5 GTPase family members.

**Fig 12.**
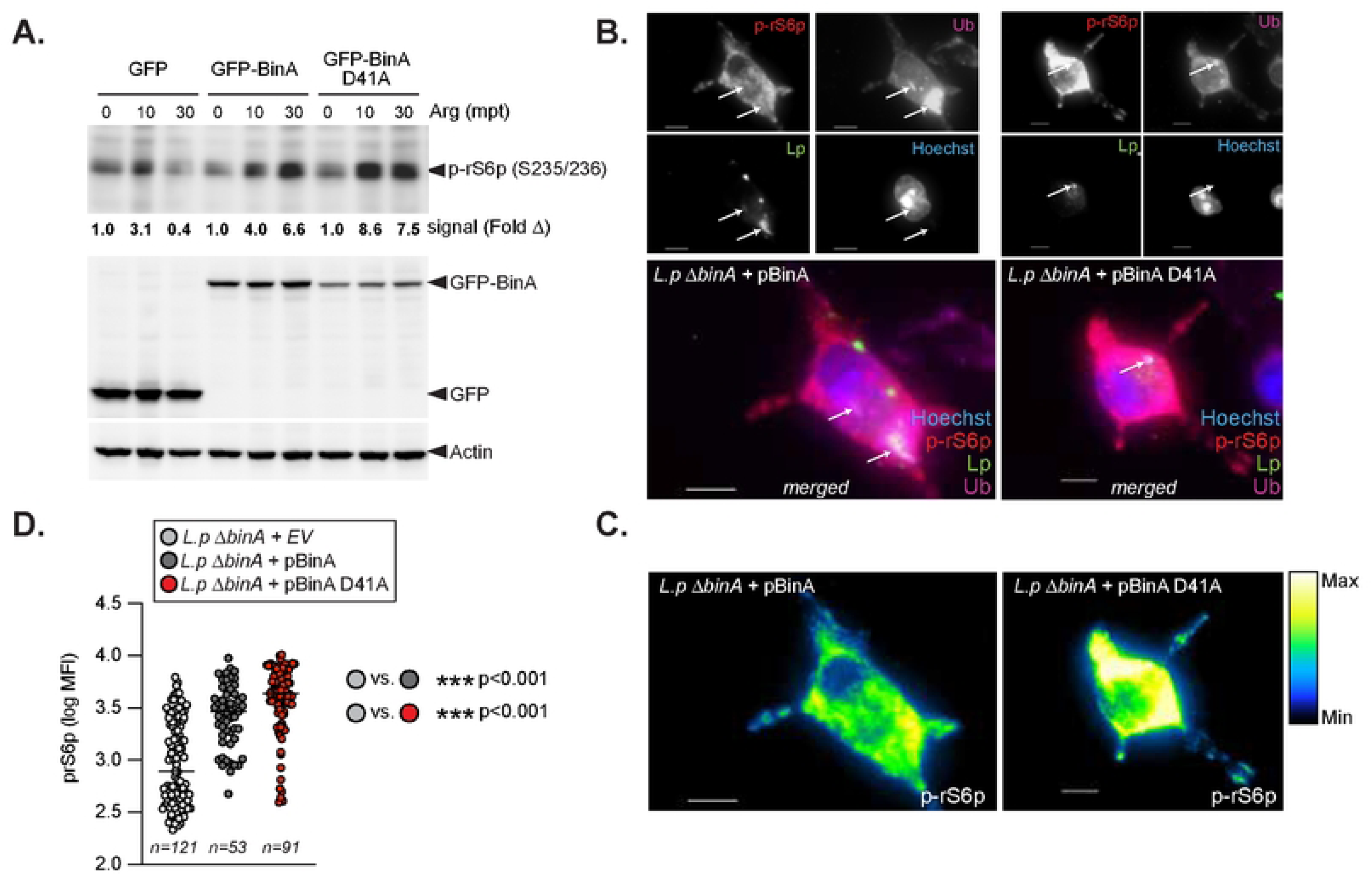
GEF activity is dispensable for TORC1 sensitization by BinA. **(A)** Kinetics of TORC1 activation triggered by starvation/refeeding stimulation with 500µM Arg in HK293 cells producing GFP, GFP-BinA or GFP-BinA D41A alleles. Band signal intensity in the phospho-immunoblots for rS6p for each condition was quantified and is presented below the figure panel as fold change from untreated cells. **(B)** Representative micrographs of TORC1 activation at 6hpi in Myd88KO iBMDMs infected by Lp01 Δ*binA* strains complemented with plasmid borne BinA or BinA D41A alleles. Micrographs from merged and their respective individual channel images are shown, where arrows indicate remodeled bacteria-occupied vacuoles decorated with ubiquitin. Phosphorylation of rS6p was used as a readout for TORC1 signaling and is shown as pseudocolor in **(C)**. **(D)** Quantitative microscopy of the p-rS6p signal at 6hpi from individual Myd88KO iBMDMs infected by Lp01 Δ*binA* strains complemented with empty vector (EV) or with the indicated plasmid borne BinA alleles. Log_10_ transformed data of p-rS6p mean fluorescent intensity (MFI) is shown where each circle represents data from a single cell. The number for cells analyzed for each condition is noted and the mean for each condition is indicated with a horizontal bar. Statistical analysis was performed with multiple comparisons Kruskal- Wallis one-way ANOVA test. The respective p- values across the different comparisons are shown. **(A-D)** One representative of three biological replicates is shown.

## Discussion

The eukaryotic checkpoint kinase mTOR is one gateway to host metabolism exploited by *Legionella* for the sustained production of membrane lipids needed for expansion of its intracellular niche – a process that benefits the pathogen by maximizing the bacterial progeny output from each infected cell. Here, we provide mechanistic insight into TORC1 subversion by *Legionella* and build upon the established model in the field, where an increase in the cytosolic amino acid pool brought by inhibition of host protein synthesis in infected cells triggers robust TORC1 signaling even in the absence of growth factor signaling [4, 48] (Fig 13A). We uncovered that the T4SS effector BinA functions as TORC1 regulator by specifically sensitizing signaling downstream of arginine and leucine stimulation through manipulation of host amino acid transport pathway via the host endosomal regulators Rab21 and Rab22. Additionally, our work demonstrates an interdependency between the cytosolic and the extracellular/intraorganellar amino acid pool for sustained TORC1 induction by *Legionella*. Several lines of evidence support the proposed model: (i) on average macrophage infections with the *binA* clean deletion mutant produced lower p-rS6p signal as compared to the parental or the complemented strains indicative of a TORC1 signaling defect; (ii) the *binA*− strain exhibited a defect in LCV expansion strictly in the presence of mTOR indicating both factors integrate in the same pathway; (iii) MTOR knockout produced a more severe LCV expansion defect as compared with the loss of *binA* consistent with the partial loss of TORC1 signaling in infections with the *binA*− mutant; (iv) Amino acid import by the Leu permease LAT-1 was reduced in macrophages infected with the *binA*− mutant and BinA was sufficient to sensitize TORC1 specifically in response to Leu and Arg; (v) BinA-induced TORC1 sensitization was lost upon stimulation with a membrane permeable TORC1 agonist consistent with BinA-dependent regulation of amino acid transport pathways. (vi) bacteria-induced TORC1 signaling during infection was rapidly attenuated upon withdrawal of extracellular amino acids and BinA expression was sufficient to prolong cycloheximide-induced TORC1 signaling in eukaryotic cells. Remains to be determined which non-cytosolic amino acid pool is tapped by BinA for TORC1 sensitization.

**Fig 13.**
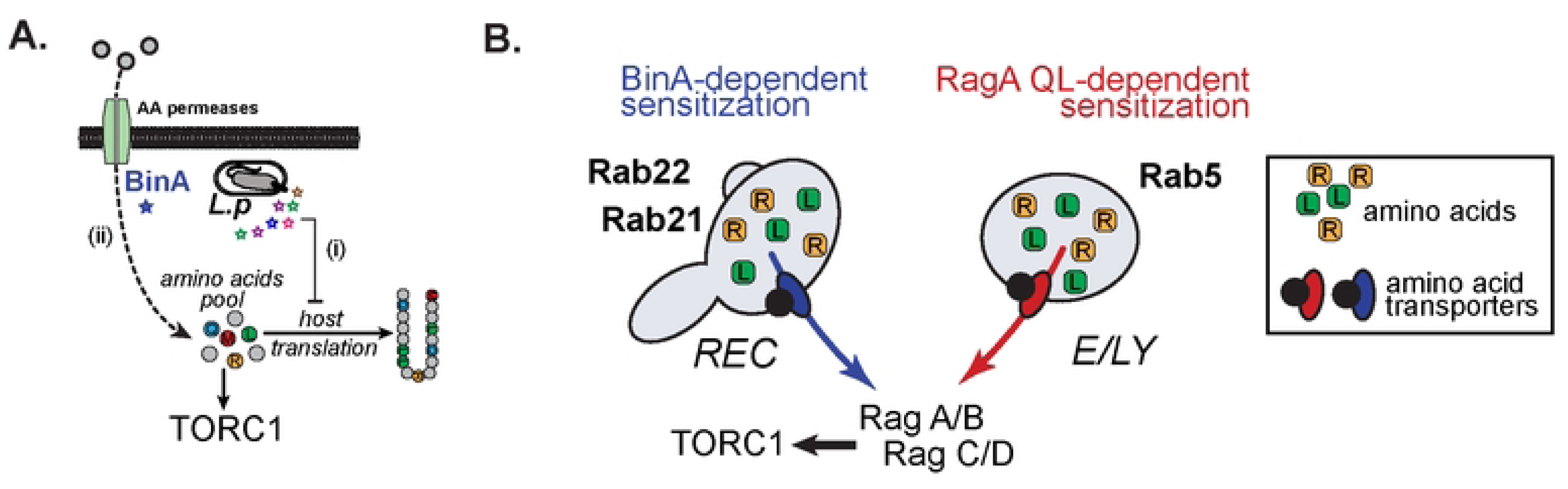
Model for BinA-dependent subversion of TORC1 activation by *Legionella*. **(A)** Early translocation of multiple T4bSS effectors causes (i) inhibition of the host translation machinery resulting in amino acids accumulation, which in turn triggers TORC1 signaling at approximately 2 hours post bacterial internalization. Enhancement of amino acid import driven by BinA (ii) sustains TORC1 signaling for several hours and represents an additional regulatory layer. **(B)** At least two distinct amino acid pools are sourced independently to provide sensory input to TORC1. One input likely originates from an organelle controlled by the small GTPases Rab21 and Rab22 such as the recycling endosomal compartment (REC). The second is regulated the Rab5 and thus is likely further down the endocytic pathway presumably of endosome/lysosome origin (E/LY). In cells containing BinA, TORC1 is sensitized through increased amino acid import from the Rab21/Rab22 compartment in a manner that also renders the Rab5-dependent pathway superfluous potentially through a switch in the utilization of transporters and/or amino acid sensors. Ectopic expression of RagA Q66L mutant elicits a preference for the Rab5- dependent amino acid pool. Both pools autonomously can trigger TORC1 signaling in the absence of BinA and RagA Q66L, which ensures TORC1 response capacity is retained irrespective of endocytic trafficking.

A large number of amino acid transporters function as heterodimers between a common CD98 heavy chain (CD98hc) that regulates trafficking and a light chain with distinct substrate specificity that function as a permease [78]. Plasma membrane CD98hc heterodimers are well-established CIE cargos that cycle through the recycling endosomal compartment [79] – a process regulated at different stages by Rab21 [73] and Rab22 [74, 80]. The BinA-dependent TORC1 sensitization is blocked by GDP-locked mutant alleles of Rab21 and Rab22 consistent with a regulatory activity localized in the early endocytic compartment (including the plasma membrane, early endosomes or recycling endosomes), which also coincides spatially with multiple amino acid transporters. However, BinA unlikely causes a pleiotropic influx of amino acids because TORC1 sensitization downstream of BinA is specific for Arg/Leu and not Met/Gln stimulation despite the capacity to all four amino acids to trigger TORC1 signaling when supplied exogenously in our assays.

The unexpected determinants switch for TORC1 signaling downstream of the Arg and Leu sufficiency responses from Rab5b (in the presence of RagA Q66L) to Rab21/Rab22 (in the presence of BinA) uncovered a novel layer of regulation in the TORC1 pathway (Fig 13B). Unlike Rab21 and Rab22, Rab5 has been previously implicated as a determinant of TORC1 activation downstream of amino acid stimulation, however the precise mechanism is still unclear [77]. Potentially, BinA might be redeploying amino acid transporters between different organelles or causing accumulation of specific intraorganellar amino acid pools via manipulation of vesicular trafficking or switching the source of Arg/Leu cytosolic import among distinct intraorganellar pools. Indeed, Rab21 has been shown to bind LAT-1 and loss of Rab21 causes missorting of certain direct CIE cargos (LAT-1, CD44 and Basigin) from the tubular endosomal compartment to lysosomes [73]; whereas, Rab22a mediates trafficking to the tubular endosomal compartment of specific CIE cargos (CD98hc, CD147) and knock down of Rab22a blocks recycling and accumulates those cargos in the early EEA+ endosomes [74, 80]. Given our results, TORC1 sensitization by BinA in response to Leu and Arg likely impinge on regulation of protein sorting at the tubular endosomal compartment. The rapid destabilization specifically of Rab22a when bound by GDP upon withdrawal of exogenous amino acids indicates an intriguing possibility of a bidirectional relationship where amino acid sufficiency and Rab22a function are interlinked.

BinA GEF activity towards Rab5 GTPase family members is dispensable and perhaps even hinders to an extent TORC1 sensitization based on the data obtained with the D41A mutant indicating that BinA has two distinct activities. While TORC1 sensitization by BinA promotes LCV expansion, it is yet unclear how BinA GEF activity benefits *Legionella* survival in the host cell. Rab5a and Rab21 are recruited to 70% of early LCVs (at 1hpi) in RAW264.7 murine macrophage cells in a T4bSS-dependent manner [81]. It is possible LCV retention of those Rabs directly promote niche biogenesis or perhaps is caused by T4bSS effector-mediated blockage in phagosomal maturation. The second notion is supported by data showing enhanced bacterial replication upon Rab5a and Rab21 depletion in A549 type II pneumocytes cell line [81] indicating that disruption of endocytic maturation benefits intracellular replication. Likely Rab21 function initially is detrimental while the bacteria transverses the endocytic compartment, but it is subsequently beneficial for the expansion of the ER-like LCV later on through the BinA- TORC1 axis. The TORC1 activation kinetics in infected cells fits well with this notion.

How does *Legionella* maintain sufficiently large cytosolic amino acid pool to sustain TORC1 activity given its consumption of host amino acids for bacterial replication? TORC1 signaling persists over several hours and coincides with bacterial replication [4]. Inflow of amino acids from the extracellular space directly or indirectly through an intraorganellar pool replenished by exogenous amino acids provides one answer supported by our data [70, 71]. Withdrawal of amino acids rapidly extinguished TORC1 signaling under conditions that block host translation either artificially through a pharmacological inhibitor or during infection indicating that without sustained inflow of amino acids the cytosolic pool falls rapidly below the threshold needed for TORC1 activation. However, amino acids-free medium still supports limited intracellular bacterial replication [36] indicating that TORC1 signaling is more sensitive to amino acid depletion as compared to bacterial replication.

In conclusion, our work identified a novel function of the *Legionella* T4SS effector protein BinA as a sensitizer of TORC1 signaling and provided mechanistic insight into this process mediated by the host membrane trafficking regulators Rab21 and Rab22.

## Material and Methods

### Reagents

The pharmacological agents used in the study were obtained from the following suppliers - Cayman Chemical Company (Torin2); Alfa Aesar (Leucine, Arginine, Glutamine, Cysteine, and iron nitrate); Cayman Chemicals (L-Leucyl-L-Leucine methyl ester). Primary antibodies were purchased from: (i) Cell Signaling – α-Akt (pan) (C67E7), α-prS6p (S235/236, D57.2.2E), mTOR (7C10), α-pS6K1 (T389, 108D12), p70 S6Kinase (E8K6T) and rS6p (5G10); (ii) Santa Cruz - α-Actin (C-2); (iii) Cayman Chemicals - α-multiubiquitin chain conoclonal antibody (Clone FK2). The following antibody was custom produced by Cocalico Biologicals against formalin-killed bacterial strains– α-*L*. *pneumophila* (chicken IgY). Secondary antibodies and Hoechst 33342 (cat #H3570) were purchased from ThermoFisher Scientific as well as α-mouse-Alexa488 (cat #A11001), α-mouse-Alexa647 (cat #A21235), α-rabbit-Alexa647 (cat #A21244), α-chicken-IgY-FITC (cat #A16055), α- chicken-IgY-TRITC (cat #A16059).

### Bacterial strains

All *L. pneumophila* strain used in this study are detailed in Table S2 and were derived from the *L. pneumophila* serogroup 1, strain Lp01 [82] and have a clean deletion of the *flaA* gene to avoid NLRC4-mediated pyroptotic cell death response triggered by flagellin when BMMs from C57BL/6J mice are infected with flagellated *Legionella*.

#### Bacterial culture conditions

*Legionella* strains were cultured either on charcoal yeast extract (CYE) plates [1% yeast extract, 1%*N*-(2-acetamido)-2-aminoethanesulphonic acid (ACES; pH6.9), 3.3mM l-cysteine, 0.33mM Fe(NO_3_)_3_, 1.5% bacto-agar, 0.2% activated charcoal] or in complete ACES-buffered yeast extract (AYE) broth (10mg/ml ACES: pH6.9, 10mg/ml yeast extract, 400mg/l L-cysteine, 135mg/l Fe(NO_3_)_3_) supplemented with 100µg/ml streptomycin. As needed, the media were supplemented with the following: 25 μg/ml kanamycin (Kam), 100 μg/ml streptomycin, 10 μg/ml chloramphenicol, or 1 mM Isopropyl β-D-1-thiogalactopyranoside (IPTG).

#### Bacterial culture conditions for inoculum preparation

For all infection experiments, *Legionella* strains obtained from dense patches grown on CYE plates for two days were resuspended to OD_600_ of 0.5 U in 1 mL AYE, placed in 15-ml glass culture tubes, and cultivated aerobically 24 to 26 hours with continuous shaking (175 rpm) at 37°C until early stationary phase was reached (OD_600_ range of 2.0 to 3.0 U). The AYE broth cultures of Lp01 GFP+ strains harboring a plasmid carrying the *gfp* gene under the control of the P_tac_ contained 10μg/ml chloramphenicol and were supplemented with 0.1 mM IPTG between 18 to 20 h post seeding in AYE broth.

#### Inactivation of Lp genes via clean deletion (CD) mutagenesis

The Lp01 Δ*binA* allele was generated by fusing ∼1kb regions upstream and downstream of the *binA* gene. To this end the upstream region was PCR amplified from Lp01 genomic DNA using the primers UpF_CD_BamHI_*binA* and UpR_CD_EcoRI_*binA*. The downstream region was similarly produced with the primers DownF_CD_EcoRI_*binA* and DownR_CD_SacI_*binA*. The resulting fragments were linked by an EcoRI site and cloned into the BamHI/SacI sites of the gene replacement vector pSR47s creating pSR47s-CDLpg0393. The pSR47s-CD was introduced in the Lp01 Δ*flaA* strain via tri-parental mating for the generation of the Δ*flaA* Δ*binA* strain by allelic exchange of *binA* with Δ*binA* via double homologous recombination. The allelic exchange was confirmed by genotyping clonal isolates with the primers SP1_CD_*binA*, SP2_CD_*binA* SP3_CD_*binA* in a single PCR reaction that produces either a fragment of 324nt for *binA* or a fragment of 525nt for Δ*binA*. Detailed information on the primers used in this study is included in Table S3.

#### Generation of plasmid-bearing Lp strains

Plasmids were introduced in an electrocompetent Lp01 Δ*flaA* Δ*binA* strain via electroporation (voltage-1800, capacitance-25µF and resistance- 200ohms). Plasmid bearing clones were isolated after growth selection on CYE plates supplemented with 10μg/ml chloramphenicol after recovery in AYE broth cultures for 6 hours. The GFP expressing strain was generated by introduction of pAM239 plasmid. Production of BinA or BinA D41A was restored by electroporation of clean deletion *binA* strain with the pJB1806-3XFlag-BinA or the pJB1806-3XFlag-BinA D41A plasmid respectively. The pJB1806-3XFlag plasmid was introduced to generate the control Lp01 Δ*flaA ΔbinA +* EV strain.

### Construction of plasmid-borne alleles

Detailed information on the primers and plasmids used in this study is included in Table S3 and Table S4 respectively. A DNA fragment containing the *binA* gene flanked by BamHI and SbFI sites was produced in a high-fidelity PCR reaction with the primers F_BamHI_+2_*lpg0393*-*binA* and R_SbFI_*lpg0393*-*binA* using Lp01 gDNA as a template. The BamHI-*binA*-SbFI amplicon was cloned in pJB1806-3XFlag (BamHI/PstI) for bacterial expression and in pEGFP-C2 (BglII/PstI) for expression in eukaryotic cells.

The BinA D41A allele was generated via Quick Change Point mutagenesis using pEGFP- BinA as a template with the primers QC_F_lpg0393_D41A and QC_R_lpg0393_D41A. Colony PCR screen was used to identify clones carrying the D41A mutation with the QC_FSP_lpg0393_D41A and the R_SbFI_*lpg0393*-*binA* primers. Putative D41A mutant alleles were sequenced. The BinA D41A allele was PCR amplified with the primers F_BamHI_+2_*lpg0393*-*binA* and R_SbFI_*lpg0393*-*binA* and cloned in pJB1806-3XFlag (BamHI/PstI) to generate the pJB1806-3XFlag-BinA D41A plasmid.

The human alleles Rab5b, Rab5b Q79L, Rab5b S34N, Rab22a, Rab22a Q64L, Rab22a S19N, RagA Q66L, and RagA T21N were synthesized as gBlocks with flanking BglII/EcoRI restriction sites and were cloned in pEGFP-C2 plasmid digested with BglII/EcoRI. All alleles were confirmed by sequencing. The murine Rab21, Rab21 Q76L and Rab21 T31N alleles were a gift from Johanna Ivaska (Addgene plasmids # 83421, 83422, 83423; RRID:Addgene_83421, RRID:Addgene_83422, RRID:Addgene_83423) [83] .

### Murine strains and generation of primary murine bone marrow-derived macrophages

C57BL/6J *Myd88*^-/-^ mice (cat #009088) were purchased from Jackson Laboratories and housed at LSUH-Shreveport animal facility. Ethical approval for animal procedures for experiments in this study was granted by the Institutional Animal Care and Use Committee at LSUHSC-Shreveport (protocol# P-15-026). Primary BMMs were derived from bone marrow progenitors isolated from C57BL/6J mice were cultured on 10 cm petri dishes in RPMI 1640 with L-glutamine (BI Biologics, cat #01-100-1A) supplemented with 10% v/v FBS (Atlas Biologics, cat #FS-0500-AD), conditioned medium from L929 fibroblast cells (ATCC, CCL-1) (30% final volume) and penicillin/streptomycin (Thermo Scientific, cat#15140163) at 37°C with 5% CO_2_. Additional media was added at days 3, 5 and 7 from the start of differentiation. Macrophages were collected and seeded for infections on day 9.

### Cell lines derivation and culture conditions

#### Derivation of immortalized Myd88 KO BMDMs

Bone marrow cells from C57BL/6J *Myd88*^-/-^ mice were differentiated into macrophages with L-929 conditioned (30% v/v) complete DMEM medium (High glucose DMEM with L-glutamine and sodium pyruvate (Thermo Cat. 11995073); 10% hiFBS, supplemented with sodium pyruvate to 2mM final concentration; supplemented with L-glutamine to 6mM final concentration 5.5 ml of 100X L-glut; 1X Penicillin/Streptomycin) for 7 days and were immortalized with the J2 murine retrovirus. J2 producing cells were a kind gift from Dr. Jonathan Kagan (Harvard Medical School) [84]. The J2 virus-containing complete DMEM medium was produced by collecting the culture medium from J2 producing cells that were cultured for 24 hours after reaching 100% confluency. The virus-containing medium was centrifuged (400x *g*, 5 min) and the supernatant was passed through a 0.2µm filter. BMDMs were transduced with J2 virus-containing supernatant fortified with 30% L-929 conditioned medium on consecutive days. Twelve hours after the second round of transduction the medium was replaced with complete DMEM containing L-929 conditioned medium (30% v/v). For the completion of the immortalization process BMDMs were split weekly (1:2 or 1:3) and the amount of L- 929 conditioned medium was progressively reduced after every split (from 30% ® 25% → 20% → 15% → 10% → 5% → 4% → 3% → 2% → 1% → 0%). Once weaned from L- 929 conditioned medium iBMDMs were cultured in complete DMEM medium.

*Generation of* MTOR*^-/-^ U937 cell line*: U937 cell line stably expressing Cas9 were previously described [57]. Lentiviral plasmids containing guide RNAs targeting human MTOR gRNA1 (BRDN0001145082), gRNA2 (BRDN0001147813), gRNA3 (BRDN0001145868) were a gift from John Doench & David Root (Addgene plasmids # 76782, #76783, #76784; RRID:Addgene_76782; RRID:Addgene_76783; RRID:Addgene_76784) [85]. HEK293E cells were used for lentivirus production. Briefly, HEK293E cells were plated in DMEM (10% hiFBS) at confluency in six-well plates one day and 24 hours later were transfected using X-tremeGENE HP DNA transfection reagent transfection reagent according to manufacturer protocol with the retroviral envelope plasmids (pPAX2 and pCMV-VSVG) and 1µg of the gRNA lentiviral plasmid. Media was collected at 24 hours post-transfection and fresh DMEM (30% hiFBS) was added for additional 24 hours. The collected supernatants were combined, centrifuged at 500x *g* to remove cellular debris and were filtered through 0.45µm filter before they were used for transduction. Cas9-expressing U937 monocytes were plated at 0.7×10^5^ in 6-well plates and were transduced with combined supernatants from all three guide RNAs in DMEM containing 0.5µg/ml polybrene and 250mM HEPES buffer. Plates were centrifuged at 500x *g* for 30min. After 4 hours, the transduction was repeated. At 24 hours post transduction the culture medium was replaced with fresh DMEM (10% hiFBS).

#### Cell lines culture conditions

Immortalized BMMs were propagated in complete High-Glucose DMEM supplemented with L-glutamine to 6mM final concentration (ThermoScientific, cat# BP379-100), 10% v/v heat-inactivated FBS (Atlas Biologics, cat# F-0500-DR), 2mM sodium pyruvate (Quality Biologicals, cat#116-079-721), 100 U/ml penicillin and 100 µg/ml streptomycin. HK293, HK293E and A549 (all purchased from ATCC) cell lines were propagated in DMEM supplemented with 10% v/v hiFBS, 100 U/ml penicillin and 100 µg/ml streptomycin. U937 monocytes (purchased from ATCC) were propagated in RPMI-1640 with L-glutamine (BI Biologics, cat #01-100-1A), 10% v/v hiFBS, 100 U/ml penicillin and 100 µg/ml streptomycin. For macrophage differentiation, U937 monocytes were treated with 10 ng/ml Phorbol 12-myristate 13-acetate (PMA)(Adipogen) for the first 24 hours after which the cells were cultured for additional 48 hours without PMA and antibiotics. All cell lines with cultured in a tissue-culture incubator at a temperature of 37°C in the presence of 5% CO_2_.

### Amino acid starvation/refeeding in transfected HK293 cells and iBMDMs

HEK293T cells were plated in poly-L-Lysine coated 24-well plates at 0.8×10^5^ cells/well in DMEM (10% hiFBS). After 4 hours cells were transiently transfected various plasmids (300ng total DNA/well) using X-tremeGENE HP DNA transfection reagent (Roche, cat # 06366236001) according with manufacturer protocol, the medium was replaced after 48 hours and treatments were carried out at 72 hours post-transfection. The starvation/refeeding treatments in experiments with HK293 were performed as follows: (i) DMEM medium was replaced with the amino acid-free Hank’s balance salt solution (HBSS, Quality Biological, cat # 119-065-101); (ii) after 4 hour of amino acid starvation, cells were stimulated with different amino acids or LLoMe diluted in HBSS for the indicated time periods; (iii) cell were then lysed and processed for immunoblot analysis. For iBMDM treatments, 1×10^6^ cells/well were seeded in 6-well plates and were cultured in serum free (SF) RPMI overnight. Next, macrophages were infected with Lp01 Δ*flaA* bacteria in SF-RPMI (MOI=15). After 4 hours, cells were washed 2X with pre-warmed PBS and HBSS (amino acids −) or SF-RPMI (amino acids +) was added to the cells for the indicated time periods.

### Immunoblot analysis

For cell lysis preparation, culture medium was removed, cells were washed with ice-cold PBS and scrapped in complete RIPA buffer (50mM Tris HCl, pH=8.0 with 150mM NaCl, 1% Igepal CA-630 (NP40), 0.5% sodium deoxycholate and 0.1% sodium dodecyl sulfate (SDS)) containing protease and phosphatase inhibitors (Thermo Scientific, cat #A3296). The total protein concentrations for each lysate was measured with a bicinchoninic acid (BCA) protein assay (Thermo Scientific, cat #23225) and 20µg were resolved by SDS-PAGE, transferred onto nitrocellulose membrane, blocked with 5% milk and incubated overnight with primary antibodies at 4^0^C. Secondary horseradish peroxidase-conjugated antibodies were used at 1:5000 for 2h at room temperature. Chemiluminescence signals were detected and digitalized with GE Amersham imager 680 instrument and images were analyzed with ImageJ version 1.53t (NIH). For each experiment, samples were run in parallel duplicates, where one membrane was immunoblotted with phospho-specific antibodies and the other membrane was probed with antibodies against the total protein.

### Immunofluorescence analysis of TORC1 activation in infected cells

For infections, BMDMs or iBMDMs (2.5×10^5^ per well) were seed on cover slips in complete DMEM for 8 hours prior to serum starvation. For all experiments cell were serum starved for 10 hours by culturing in RPMI 1640 with L-glutamine (SF-RPMI). Cell were infected with *Legionella* at MOIs as indicated for each experiment and at 60 min post infection extracellular bacteria were removed by washing 5X with pre-warm PBS. Cells were cultured in SF-RPMI for the duration of the infection.

For immunostaining, coverslips containing cells were washed with 3x PBS and fixed with 2% paraformaldehyde for 20 min, permeabilized with cold methanol for 30 seconds, blocked with 2% BSA in PBS for 60 min. Primary antibodies dilutions were 1:500 for α-p- rS6p, 1:200 for α-ubiquitin (FK2), and 1:1000 for all α-*Legionella* antibodies. Primary antibodies were incubated in PBS containing 1% BSA overnight at 4°C. Secondary antibodies were used at 1:500 dilution and Hoechst 33342 at 1:2000 for 60 min at room temperature. Coverslips were mounted with ProLong Gold antifade reagent (ThermoFisher) onto slides and examined by fluorescence microscopy.

### Microscopy analyses of infected cells

Images were acquired with inverted wide-field microscope (Nikon Eclipse T*i*) controlled by NES Elements v4.3 imaging software (Nikon) using a 60X/1.40 oil objective (Nikon Plan Apo λ), LED illumination (Lumencor) and CoolSNAP MYO CCD camera. Image acquisition and analysis was completed with NES Elements v4.3 imaging software. Only linear image corrections in brightness or contrast were completed. For all analyses, three-dimensional images of randomly selected fields were acquired, and image acquisition parameters were kept constant for all the cover slips from the same experiment. For phospho-rS6p analysis in cells, the ubiquitin signal was utilized to define a binary mask for each cell from which the mean fluorescence intensity of p-rS6p was obtained. Background fluorescence signal from each acquired field was individually determined from a cell-size mask positioned in an unoccupied area and subsequently subtracted from the MFI of each cell in the field.

### Automated time-lapse live-cell imaging of *Legionella* growth with host cells

U937 monocytes were seeded at 1×10^5^ cells/well in 96-well black-wall clear-bottom plates (Corning, cat# 3904) and differentiated into macrophages in RPMI media supplemented with PMA for 24 hours followed by 48 hours culture in PMA-free RPMI medium. All infections were carried out at MOI = 2.5 with AYE broth grown bacteria in the presence of 1mM IPTG. Final media volume was 150µl per well. In each experiment, all conditions were performed in technical triplicates. Plates were centrifuged (500 rpms, 5min) and were loaded into the IncuCyte™ S3 microscope housing module. The IncuCyte™ S3 HD live-cell imaging platform (Sartorius) is a wide-filed microscope mounted inside a tissue culture incubator and is controlled by IncuCyte™ software. For each well, four single plane images in bright field and green channel (ExW 440-480nm/EmW 504-544nm) were automatically acquired with *S* Plan Fluor 20X/0.45 objective every four hours over several days. Images were analyzed with the IncuCyte™ Analysis software. For quantitation of bacterial replication, GFP signal was used for the generation of a binary mask that defined total signal integrated intensity (GCUxµm^2^) in each image or for each GFP+ object when LCV expansion was measured. For LCV analysis, bacterial fluorescence was used to generate a binary mask to define individual LCV objects and to measure the object’s area size (GCUxµm^2^) and the number of objects per image. For single object LCV tracking, the LCV area was recorded for ∼100 LCVs per condition for every timeframe from appearance of the LCV to its disappearance. The Incucyte S3 imaging analysis was performed in the Innovative North Louisiana Experimental Therapeutics program (INLET) core facility at LSU Health-Shreveport (RRID: SCR_024990).

### Loss-of-function screen for TORC1 regulating Lp T4bSS effectors

Arrayed effector mutant transposon library used in this study has been previously described in detail [52]. TORC1 activation in primary MyD88 KO BMDMs infected by Lp was analyzed via immunofluorescence for p-rS6p (Ser235/236) as detailed in the ‘*Immunofluorescence analysis of TORC1 activation in infected cells*’ methods section. To this end, Myd88 KO BMDMs were seeded at 2.5×10^5^ cells/well on 1.5mm glass coverslips placed in each well of a 24-well plate in replating medium for 8 hours prior to serum starvation. For all experiments cell were serum starved for 16 hours by culturing in RPMI 1640 with L-glutamine (SF-RPMI). Transposon mutants were grown on CYE agar plates for 48 hours from which liquid cultures in AYE medium were grown overnight as detailed in the ‘*Bacterial culture conditions for inoculum preparation*’ methods section. Infections were performed in technical triplicates for bacterial strain (∼ MOI=15) and in each set of infections a wild-type control strain (Lp01 Δ*flaA*) as well as uninfected macrophages were used to provide baselines for TORC1 signaling. All infections were synchronized at 2hpi and were stopped at 8hpi.

Imaging of immunolabelled infected macrophages was completed as detailed in the ‘*Microscopy analyses of infected cells*’ methods section. For each infection the percentage of infected macrophages with moderate/high p-rS6p signal (MFI > 400) was scored taking into account LCV remodeling (i.e. presence of Ub signal at the LCV) as an additional criterion. More than 20 mutant strains were analyzed in each experimental set of infections, where the mean percentage of p-rS6p+ infected cells as well as the standard deviation of the data was calculated by averaging the data from all the bacterial strains in the set. The z-score (standard deviations away from the mean) was calculated for each bacterial strain. Data is provided in Table S1.

### Assay of boronophenylalanine (BPA) uptake via LAT1-mediated transport

BPA - a phenylalanine derivative - is internalized via the amino acid transporter LAT-1. BPA uptake assay (Dojindo Labratories, cat# UP04) was used to measure BPA transport in Lp-infected macrophages as per the suggested manufacturer’s protocol. Briefly, 1×10^5^ Myd88 KO iBMDMs were seeded in each well of a 96-well black wall microplate and were cultured at 37°C overnight in a 5% CO2 incubator in complete DMEM medium. Macrophages were infected in SF-RPMI medium with the indicated strains grown in overnight AYE liquid cultures at MOI = 20. Infections were synchronized at 2 hpi by several washing cycles with pre-warmed PBS. At 4 hpi, the SF-RPMI culture medium was removed and the cells were washed three times with pre-warmed HBSS. Next, pre-warmed HBSS containing BPA was added to each well and macrophages were allowed to accumulate L-BPA for 5 min in a tissue culture incubator at 37℃. BPA was omitted from the ‘blank’ control. Next, culture supernatants were removed, and cell were washed three times with pre-warmed HBSS, after which HBSS containing the BPA-specific fluorescent probe was added to each well for 5 minutes. LAT-1 transport activity was inhibited with 1 mM 2-aminobicyclo [2.2.1]-heptane-2 carboxylic acid (BCH), which was added 10 min prior to L-BPA addition and was maintained for the duration of the experiment. Fluorescence was measured with a Tecan Spark plate reader using the DAPI filter (*Excitation* 360±30nm/ *Emission* 480±30nm) and the gain was set at 75. For each well, the data represents the mean fluorescent signal from 25 tiled area scans that cover the entire surface of the well. Immediately after a plate was imaged the cell were lysed with RIPA cell lysis buffer and the total protein content was measured against a BSA standard. Every experimental condition was run in four technical replicates and the data is presented as fluorescence units/mg protein.

## Acknowledgements

We would like to thank the INLET high-throughput imaging core at LSUHSC-Shreveport (RRID: SCR_024990) for technical assistance.

## Supporting information

**S1 Figure. Sensitization of leucine-dependent TORC1 signaling by BinA is regulated by Rab21 and Rab22a but not Rab5.** Kinetics of TORC1 activation triggered by starvation/refeeding stimulation with 100µM Leu in HK293 cells producing BinA in the presence or absence of different Rab5 **(A-B)**, Rab21 **(C-D)**, or Rab22a **(E-F)** alleles. **(B, D and F)** Quantitative analyses of band signal intensities show Averages ± StDev from at least three biological repeats (left panels) and the respective area-under-the curve (AUC) analyses are presented in the right panels. Statistical analyses were completed with one-way ANOVA with Dunnett’s multiple comparison test using the ‘BinA+GFP’ as control group and p-values are indicated in the respective data panels.

**S2 Figure. Rab5 family GTPs are dispensable for TORC1 signaling triggered by leucine in the absence of BinA or RagA Q66L sensitization.** Kinetics of TORC1 activation triggered by starvation/refeeding stimulation with 400µM Leu in HK293 cells producing GFP or different Rab5 **(A)**, Rab21 **(B)**, or Rab22a **(C)** alleles. Band signal intensity in the phospho-immunoblot for rS6p for each condition was quantified and is presented below the respective figure panel as fold change from untreated cells. The data shown is from one experiment out of three biological replicates.

**S3 Figure. Withdrawal of extracellular amino acids reduces the abundance specifically of the GDP-locked Rab22a S19N allele.** HK293 cells ectopically producing GFP alone or the indicated GFP-tagged Rab22a alleles were treated with HBSS for 4 hours. Immunoblot analysis demonstrates reduction of total Rab22a S19N abundance upon amino acid withdrawal. Band signal intensity for GFP blots for each condition was quantified and is presented below the respective figure panels as percentage of signal from cells that were not treated with HBSS. The data shown is from one experiment out of three biological replicates.

**S4 Figure. Ectopic production of BinA but not BinA D41A enlarges Rab5-containing endosomes. (A-B)** Representative micrographs of A549 cells co-expressing RFP-tagged Rab5 and either GFP-BinA **(A)** or GFP-BinA D41A **(B).** Merged and the respective single channel images are shown. **(C)** Quantitative analysis of large Rab5 vesicles produced by BinA and BinA D41A-expressing cells. The graph shows Averages ± StDev from three biological replicates where at least 100 cells were counted for each condition. Statistical analysis was completed with unpaired T-test with Welch’s correction and the p-value is indicated in the graph panel.

**S1 Table. Forward genetic transposon-based screen for bacterial TORC1 regulators.** List of all transposon mutants included in the forward genetic screen and their respective z-scores.

**S2 Table. Bacterial strains used in this study.**

**S3 Table. Primers used in this study.**

**S4 Table. Plasmids used in this study.**

